# The Attentional Thief: Scrolling-Like Picture Streams Compress Subjective Time

**DOI:** 10.64898/2026.07.02.734699

**Authors:** Chunyu Qu, Artyom Zinchenko, Siyi Chen, Zhuanghua Shi

## Abstract

People lose track of time while scrolling, but real feeds confound novelty, reward, social signals, and self-paced control. We asked whether a scrolling-like picture stream, advanced at the viewer’s own pace, compresses subjective time on its own. Twenty-three adults (10 female; age *M* = 27.0 years, *SD* = 4.1, range 22–36) completed three within-participant conditions with EEG and eye tracking. In Scrolling, they advanced through natural photographs by mouse click at their own pace, each preceded by a brief fixation interval. In Watching, they passively viewed new photographs with the picture and fixation durations of their own Scrolling block, permuted. In Baseline, a single photograph from the same pool stayed on screen for the matched duration. After each block, participants estimated its elapsed duration. Both sequential-picture conditions were judged far shorter than Baseline (Scrolling 48%, Watching 52%, Baseline 65% of clock time; Scrolling versus Baseline *d*_z_ = −0.80), a compression carried by the fixation intervals between pictures, which observers barely counted. Scrolling was judged shorter still than Watching (*d*_z_ = −0.77); the size of this increment varied with sequential context, in a pattern of assimilation toward the preceding estimate. EEG and eye tracking characterized self-paced viewing as a changed anticipatory state, an earlier visual response, and more active visual sampling. The results fit an attention-weighted readout of time: a picture stream and its uncounted intervals draw attention from the clock. Time processing during feed-like viewing is vulnerable to the structure of sequential viewing, independent of content or reward.

## Introduction

Social media users often report losing track of time: a quick glance at a feed stretches far past what was intended. Field studies bear the report out. Social-media episodes are marked by deep absorption and reduced awareness of elapsed time (Baughan et al., 2022); adolescents’ estimates of their own smartphone time correlate only weakly with 45 days of logged use, but the size of the misjudgment predicts problematic use a year later (Marciano & Camerini, 2022), and short-form video use distorts time perception, most among heavy or problematic users (Jiang et al., 2025; Yang et al., 2024). These studies establish the association but cannot say which feature of the platform produces it, because in everyday use, algorithmic curation, social reward, changing content, and the user’s own pacing arrive together. We isolate the most basic of these features: the scrolling-like sequence itself, a stream of discrete events, such as posts, clips, or images, separated by brief gaps and advanced at the viewer’s own pace. Isolating this feature matters for theory. The two leading accounts of prospective timing (Roseboom et al., 2019; Zakay & Block, 1997) make opposite predictions for such a sequence: compared with a single unchanging scene, one predicts that it will feel shorter, the other that it will feel longer. The direction of the effect therefore shows which process dominates. Isolating it also matters for practice. Stopping when one intended depends on keeping track of how much time has passed, so a structural feature that erodes this awareness is worth understanding regardless of the content on screen.

How long an interval feels depends on what the observer processes during it, and two accounts disagree about how. The attentional-gate account treats prospective duration as a count of pacemaker pulses that accumulate only while attention is on the clock. When competing demands pull resources toward non-temporal processing, fewer pulses are counted, and the interval feels shorter (Zakay & Block, 1997). Loading working memory or adding a concurrent task shortens estimates of intervals, whereas instructing observers to monitor time lengthens them (Polti et al., 2018; Zang et al., 2026), and prospective reproductions of everyday events shrink as the events grow richer (Bangert et al., 2019). A content-readout account builds duration from the content of the interval instead: the estimate accumulates the salient changes registered in perceptual processing, so an interval packed with distinct events is judged longer than a sparse one. Prospective estimates of natural-scene videos follow this pattern, with walks through a busy city judged longer than scenes in a cafe or office, and salient events in visual cortex activity predict trial-by-trial biases in duration reports (Roseboom et al., 2019; Sherman et al., 2022). At the scale of a second, too, intervals filled with tone sequences seem longer than empty or continuously filled ones (Horr & Di Luca, 2015). The two accounts differ on what a sequence of events should do: draw attention from time and shorten the estimate, or supply more salient changes and lengthen it.

Self-paced advance is not a single process. The scrolling gesture bundles multiple processes together: a decision that the current event has been viewed long enough, the motor act that carries it out, the resulting dwell time and near-perfect predictability of the next onset, because the viewer triggers it. Voluntary action of this kind shapes both the timing of what follows and the attention paid to it (Haggard et al., 2002), and each component could draw attention from the clock.

We compared three within-participant viewing conditions in which the events were natural photographs drawn from one category-balanced pool (Figure 1). In Scrolling, participants advanced through a sequence of photographs by mouse click at their own pace. Each picture was preceded by an inter-image fixation interval, a fixation cross on a gray field that formed the gap between pictures, serving to return gaze to the center of the screen and loosely mimicking the loading pause between items in a feed. In Watching, participants viewed new photographs without self-scrolling. Each block copied the durations of the participant’s own Scrolling block of the same length, jointly permuted, so that the two conditions shared the same set of picture durations, the same total fixation time, and the same total block time, but not the same photographs or temporal order. We refer to Scrolling and Watching together as the sequential-picture conditions. In Baseline, a single photograph from the same pool stayed on screen for the duration of the matched Scrolling block, with one fixation cross before it, so Baseline contains neither picture transitions nor fixation intervals. Blocks held 9, 12, or 18 pictures and lasted about a minute on average. After every block, participants estimated how long the block had lasted, in seconds, including fixation intervals. Because they knew from the outset that every block would be timed, the estimates are prospective. A single old/new recognition judgment followed each estimate.

**Figure 1.**
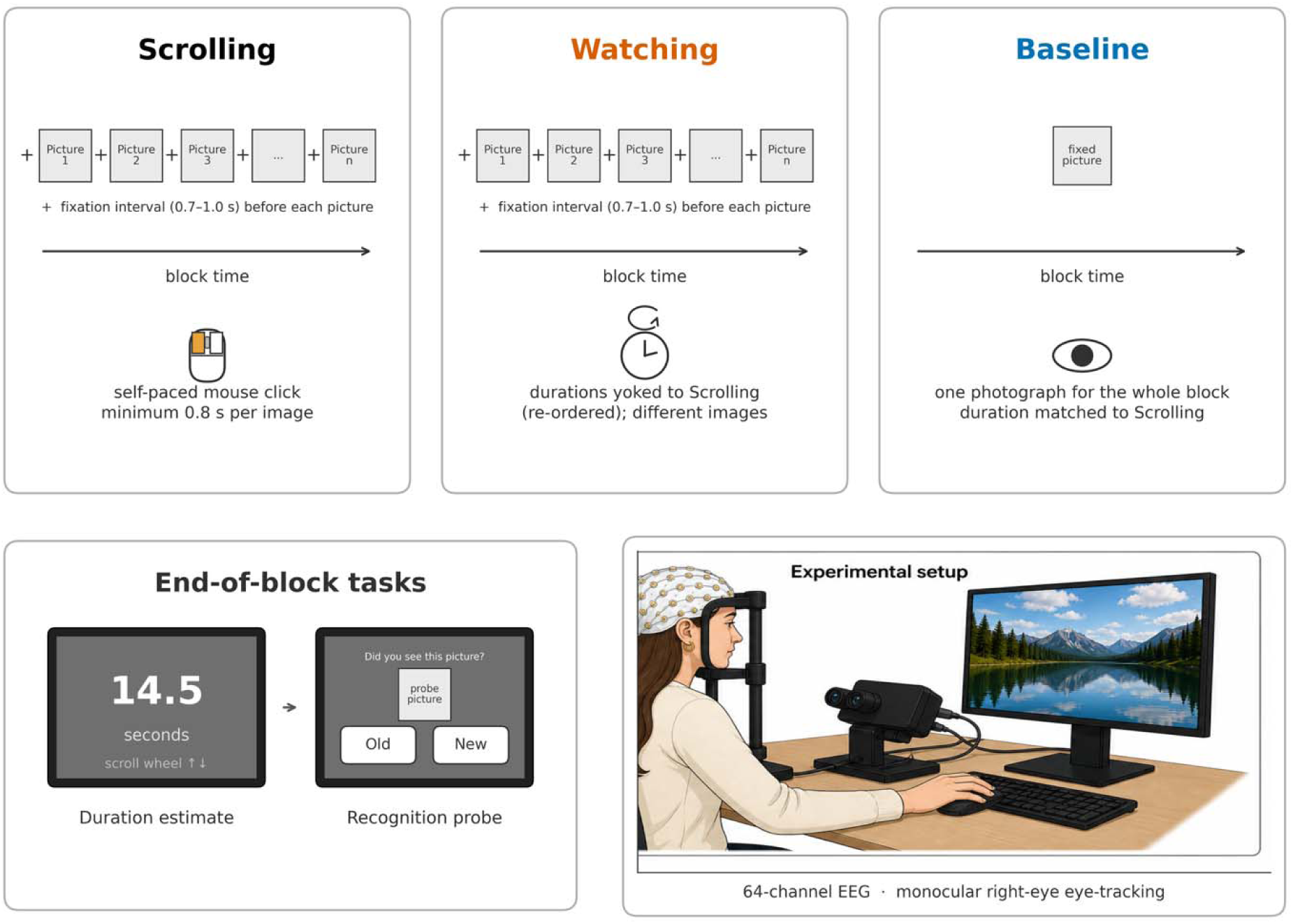
Design schematic. Three within-participant conditions, matched on total block duration within each repetition, target self-paced visual advance. Under Scrolling, participants advanced through a sequence of natural photographs at their own pace by mouse click, each picture preceded by an inter-image fixation interval. Under Watching, participants passively viewed different photographs from the same pool, presented with the picture and fixation durations of their own Scrolling block of the same repetition and block length, jointly permuted. Under Baseline, one photograph remained on screen for the whole block, with no picture transitions. After every block, participants estimated the elapsed duration of the whole block and judged whether a single probe image was old or new. Sixty-four-channel EEG and monocular right-eye eye tracking were recorded throughout. The pictures are shown schematically by text placeholders. The actual stimuli were obtained from Pexels and used in accordance with the Pexels License.

The design yields two contrasts, one for each question raised above. Comparing the sequential-picture conditions with Baseline sets a stream of events against a single unchanging scene, the comparison on which the two timing accounts diverge. Comparing Scrolling with Watching targets the self-paced bundle as a whole: Watching keeps the participant’s own duration statistics but removes the decision, the act, the dwell choice, and the predictability together. This second contrast measures what self-paced advance adds without separating these components.

If self-paced advance shortens the estimate, it could draw attention from the clock in two ways. Viewers who decide when to move on may process each picture more deeply once it appears, taking attentional resources away from time. Alternatively, because they trigger each onset themselves, they may prepare for the next picture before it arrives. EEG and eye tracking, recorded throughout, can tell these two readings apart. Picture-locked potentials at a posterior-occipital cluster capture the state of visual cortex before a picture appears and the response the picture evokes in the early P1-like, P2-like, and late positive windows (Hillyard & Anllo-Vento, 1998). Deeper processing should make the picture-evoked response larger under Scrolling than under Watching after onset, regardless of pre-onset baseline correction. Preparation should instead make the conditions differ already before onset. That difference should then carry into the baseline-corrected windows with no post-onset difference of its own, and the timing of the early response may shift. Eye movements show how actively the eyes sampled each scene, and so does the occipital potential evoked as each saccade lands (the lambda response, Madison et al., 2025). The recognition probe, one old/new judgment per block, checks that participants attended to the pictures in both sequential-picture conditions.

Four predictions follow. First, the sequential-picture conditions should be judged shorter than Baseline under the attentional-gate account, because processing a stream of eventful pictures pulls attention from the clock (Bangert et al., 2019; Polti et al., 2018; Zakay & Block, 1997), and longer than Baseline under the content-readout account, because a stream of distinct pictures carries more salient changes than one unchanging photograph (Horr & Di Luca, 2015; Roseboom et al, 2019; Sherman et al., 2022). Second, Scrolling should be judged shorter than Watching if self-paced advance draws attention from time, because deciding when to move on and executing the move consume resources that would otherwise reach the clock. Identical duration statistics in the two conditions rule out exposure time as the source. Third, the inter-image fixation intervals should be underweighted in the estimates, because a scrolling-like sequence pulls attention toward the forthcoming image (Nobre et al., 2007; Nobre & van Ede, 2023). In Scrolling, each interval also separates the viewer’s click from the picture it brings, an action-outcome interval that voluntary action subjectively compresses (Haggard et al., 2002; Wenke & Haggard, 2009). Fourth, if Scrolling is judged shorter than Watching, the electrophysiological and oculomotor record should show which reading fits: deeper processing as a larger post-onset picture response, preparation as a difference already present before onset. Recognition accuracy should not differ between the sequential-picture conditions if both were attended.

## Methods

### Participants

Twenty-seven adults took part. Four were excluded during data collection^1^. The analyzed sample comprised 23 participants (10 female, 13 male; age M = 27.0 years, SD = 4.1, range 22– 36). Age and gender were the only participant characteristics recorded; no information on race, ethnicity, nationality, socioeconomic status, educational attainment, or habitual media use was collected. Participants gave written informed consent and received monetary compensation (15 EUR/h). The study was conducted under ethical approval granted by the Ethics Board of the Department of Psychology at LMU Munich (approval date 27.09.2021), and all participants gave written informed consent.

The sample size was determined by an a priori power calculation targeting one contrast: the paired Scrolling-versus-Baseline difference in the raw duration estimate. This contrast is the closest analogue in our design to the eventful-movie versus blank-fixation contrast of Bangert et al. (2019), who found *d* = 0.64 for it. A G*Power calculation (paired *t* test, two-sided, *d* = 0.64, α = .05, 1 − β = .80) gave a minimum of 22 participants. The calculation did not cover the Scrolling-versus-Watching contrast, the repeated-measures ANOVAs, or the EEG and eye-tracking tests. With N = 23, the paired contrast has 80% power for dz ≥ 0.61. For the 3 × 3 repeated-measures ANOVA, with the observed Greenhouse–Geisser ε, the smallest effect detectable with 80% power at α = .05 is η_p_² ≈ .17 for the condition main effect and η_p_² ≈ .09 for the Condition × Block length interaction. The interaction was not a hypothesis of the study, and interactions smaller than this bound could not have been detected.

### Stimuli and apparatus

The experiment was run in PsychoPy 2025 with stimuli presented on a Samsung S22E450 monitor (47.4 × 29.6 cm active display, 1680 × 1050 pixels, 60 Hz) viewed at 57 cm (35.3 pixels/°). Stimuli were natural photographs balanced across three categories: human, social or man-made, and nature. We chose category-balanced photographs over material from a live feed because the question concerns the act of self-paced advance itself. A live feed adds personalized content, reward, and social contingencies, and algorithmic ordering that vary across participants and cannot be yoked between conditions. A fixed, category-balanced pool, by contrast, holds content variability constant across conditions and participants, and personalization and platform contingencies are absent by design. Photographs were obtained from Pexels via the official Pexels API and used in accordance with the Pexels License. Each block contained equal numbers of images from the three categories (block lengths 9, 12, and 18 pictures), no photograph was repeated anywhere in a session, and the Baseline photograph was a random category-balanced draw from the same pool. Photographs were assigned to blocks at random within category for each participant, so no photograph was tied to a condition. Images filled a 32 × 12° visual-angle window, and a central fixation cross marked transitions against a calibrated gray background (12.32 cd/m²). The mean luminance of the stimulus pool (13.58 cd/m²) closely matched the fixation background, minimizing transient luminance artifacts. As a balance check, whole-image statistics of the photographs actually presented did not differ across luminance, RMS contrast, edge density, spatial-frequency slope (Supplement S1, full table S1.1). These checks concern low-level features only; semantic complexity was not measured.

We recorded EEG using a Brain Products actiCAP system (64 active scalp electrodes, 1000 Hz sampling rate, FCz online reference, AFz ground) and monitored blink activity via VEOG. We tracked gaze monocularly from the right eye (1000 Hz) using an EyeLink 1000. Calibration involved a nine-point routine performed at the start and repeated three times mid-session. EEG and eye tracking were synchronized through shared parallel-port trigger codes and EyeLink message events emitted from the same PsychoPy process.

### Procedure

We compared three within-participant conditions, matched on total block duration within each repetition: Scrolling, Watching, and Baseline. In Scrolling, participants advanced through a sequence of natural photographs at their own pace by mouse click. Each picture remained on screen until the click (minimum 0.8 s) and was preceded by an inter-image fixation interval, a fixation cross on the gray background lasting 0.7 to 1.0 s (jittered). We refer to these as fixation intervals below. In Watching, participants passively viewed a stream of new photographs, category-balanced and drawn from the same pool, whose picture and fixation durations were copied from the participant’s own Scrolling block of the same repetition and block length and then jointly permuted: one random permutation was applied to the picture durations and the fixation durations together. The joint permutation preserves the multiset of picture and fixation durations, the total block time, and the total fixation time, and it breaks the pairing between a particular dwell decision and a particular picture. Watching is therefore a passive stream carrying the participant’s own timing statistics rather than a reproduction of the participant’s inspection sequence. It matches Scrolling in duration statistics while removing self-paced control. The two conditions also differ in picture identity, in the predictability of picture onsets, and in session position. In Baseline, one fixation interval (0.85 s on average) preceded a single photograph from the same pool, which remained on screen for the duration of the Scrolling block of the same repetition and block length. The block therefore contained no picture transitions and no further fixation intervals. Figure 1 illustrates the three conditions and the timing of the duration-estimation and recognition probes.

Four repetitions of the 3 (condition) × 3 (block length: 9, 12, or 18 pictures) design yielded 36 blocks per participant. Block lengths were shuffled within each condition’s three blocks, and a 2.5 s pause separated successive blocks. Scrolling ran first within every repetition because the Watching and Baseline blocks took their durations from it. Watching and Baseline then alternated between the second and third positions across repetitions, with the starting order randomized per participant, so that each of the two occupied each position in two of the four repetitions. Each condition’s three blocks ran as one sequence introduced by an instruction screen naming the condition, and an optional short break followed every sequence. At the end of the first three repetitions, the break included an eye-tracker recalibration. The Scrolling sequence therefore followed the practice block in the first repetition and the recalibration break in every later one.

After every block, participants estimated the elapsed duration of the whole block, including its fixation intervals, in answer to the on-screen prompt “How long was that (in seconds)?”, using a slider (range 0–120 s, 0.5 s steps with the mouse scroll wheel). Participants were told before the experiment began that they would estimate each block’s duration, so the timing judgment was prospective from the first block onward: knowing in advance that elapsed time would be queried, participants could attend to time while viewing every block, including the first. An old/new recognition probe followed (left/right mouse click).

The recognition probe presented a single image and asked whether it had appeared in the block just completed. Its role was a secondary check that participants attended to and encoded the pictures in both sequential-picture conditions, and it was not designed as a test of memory models. Old and new probes were each presented on 18 of the 36 blocks, with the old/new schedule shuffled across blocks at the start of the session. On old-probe trials, the target was drawn randomly from the just-completed block’s image set; on new-probe trials, the target came from a pre-allocated foil pool of 18 category-balanced images (six per category), each used once. Because the probe delivered one old/new trial per block, recognition accuracy in each condition rested on roughly twelve binary trials per participant (four repetitions × three block lengths).

### EEG preprocessing

EEG was preprocessed in MNE-Python (Gramfort et al., 2013). Each raw recording was loaded with its standard 10–20 montage, and the online reference (FCz) was recovered as a flat channel before re-referencing to ensure fronto-central electrodes remained recoverable. Bad channels were detected from pre-filter amplitude statistics, flagging channels flat below 0.1 µV (SD) or noisy above 200 µV (1000 µV for participant sub-003). We manually inspected TP9 and TP10 because they served as the linked-mastoid reference. In all participants, these electrodes maintained satisfactory signal quality and required neither rejection nor interpolation.

Continuous data were then bandpass filtered from 0.1–30 Hz and notch-filtered at 50 Hz (MNE defaults: zero-phase FIR, Hamming window, firwin transition bandwidth). The 0.1 Hz high-pass cutoff removes slow drift while sparing the slow late positivity that a higher cutoff would distort, and the 30 Hz low-pass attenuates muscle and line noise without altering the morphology of the posterior components analyzed here (Widmann et al., 2015). Data were re-referenced to linked mastoids (TP9 and TP10), yielding 65 EEG channels. Because free viewing produces ocular artifacts on every trial (blinks, the corneo-retinal potential of saccades, and the saccadic spike potential), we removed them with Independent Component Analysis, the established approach for coregistered eye-movement and EEG data (Dimigen et al., 2011). We fitted the ICA (FastICA, full-rank, fixed random seed, 250 µV rejection threshold) on a 1 Hz high-pass copy of the data, on which the decomposition is more stable, and used ICLabel (mne-icalabel) to classify components automatically and reproducibly (Pion-Tonachini et al., 2019), excluding those with an “Eye” probability ≥ 0.8 or other noise-category probabilities (Muscle, Heart, Line Noise, or Channel Noise) ≥ 0.9. Finally, we projected the retained components back to the 0.1 Hz filtered data and spherically interpolated any remaining bad channels.

Picture-locked epochs spanned −200 to +1000 ms around image onset (one epoch per Scrolling, Watching, or Baseline picture trigger) and were baseline-corrected using the 200 ms pre-onset window (Alday, 2019). Epochs were cleaned by AutoReject, which learns per-channel rejection thresholds by cross-validation instead of a single subjective amplitude cutoff (Jas et al., 2017), with a ±150 µV peak-to-peak fallback; participants whose rejection rate exceeded 30% were flagged for manual review and retained when condition-wise residual counts supported analysis. Fixation-locked epochs spanned −200 to +800 ms around fixation onset on the post-ICA continuous recording (no second ICA pass), with the same baseline window and a ±150 µV peak-to-peak threshold; fixations shorter than 80 ms or longer than 1500 ms were discarded before epoching to remove blink artifacts and micro-fixations. After cleaning, each participant retained on average 140 picture-locked epochs in Scrolling (range 62–159), 140 in Watching (69–156), and 10 in Baseline (6–12). In total, 10.3% of the 7,437 picture epochs were rejected, and three participants with rejection rates above 30% were retained after inspection. Fixation-locked epochs per participant averaged 1,024 in Scrolling (342–3,345), 963 in Watching (383– 2,646), and 1,058 in Baseline (418–2,484), and Supplement S2 shows their distribution across participants.

### Eye-tracking acquisition and ET–EEG synchronization

EyeLink recordings (1000 Hz, right eye) provided discrete fixation, saccade, and blink events alongside message markers for experiment metadata (block start/end, image onset/offset, and trial conditions). Eye tracking serves two roles: it describes how participants sampled the pictures under each condition (fixation rate, fixation duration, saccade amplitude), and it supplies the fixation onsets to which the fixation-related potentials are time-locked, following the coregistration approach of Dimigen et al. (2011). To synchronize the eye-tracking and EEG streams, we mapped fixation onsets onto the EEG timeline using local image-onset anchors, which yielded a median within-block root-mean-square error (RMSE) well under 20 ms.

From the EyeLink event stream, we derived per-block oculomotor summaries, including fixation rate, mean fixation duration, and saccade amplitude, converting pixel amplitudes to degrees of visual angle using the calibrated screen resolution (35.3 pixels/°). Each measure describes a different aspect of visual sampling: fixation rate (fixations per second of viewing) shows how often gaze moved to a new location, mean fixation duration how long each location was inspected, and saccade amplitude how far gaze traveled between locations. Before computing these metrics, we removed blink-transition artifacts and positions that exceeded the display bounds plus a 100-pixel margin.

### Behavioral analyses

Behavioral, ERP, and FRP analyses shared the same inferential tools, but their factorial designs differed by modality. For the timing measures, we entered participant-level means for each condition (Scrolling, Watching, Baseline) and block length (9, 12, 18 pictures) into 3 × 3 repeated-measures ANOVAs, with effect sizes (η_p_²) computed from the F-statistic. Recognition accuracy was analyzed with a 2 × 3 repeated-measures ANOVA comparing Scrolling and Watching across the same block lengths. Given that the EEG measures were based on individual picture presentations, we considered the block condition as the only within-participant factor.

The fixation-locked lambda entered a one-way, three-condition repeated-measures ANOVA, whereas the picture-locked component windows were tested with planned Scrolling-versus-Watching contrasts, as Baseline had too few trials. We assessed sphericity using Mauchly’s test and applied Greenhouse–Geisser corrections where violations occurred. Following omnibus tests, we conducted planned paired t-tests (collapsed across block lengths and Holm-corrected) to contrast the three conditions. Paired contrasts are reported with the mean difference and its 95% CI, the SD of the within-participant differences, and the within-participant correlation between the two conditions. Effect sizes for paired contrasts are reported as Cohen’s dz with 95% noncentral-t confidence intervals, except for picture-locked windowed amplitude contrasts, which used 95% bias-corrected and accelerated bootstrap CIs (10,000 resamples).

We analyzed three primary measures: estimated duration (s), estimation ratio (estimate / actual), and recognition accuracy. For the recognition Scrolling-vs-Watching contrast, we reported a two-one-sided test (TOST) of equivalence alongside the conventional contrast, setting a post hoc smallest effect size of interest (SESOI) of dz = ±0.50, the smallest effect plausibly detectable given the low-density probe (≈12 binary trials per condition).

To determine whether raw differences in duration estimates arose from neglecting the fixation intervals, we performed a post hoc accounting analysis. We defined fixation time as the sum of all fixation intervals within a block. For Baseline, we set fixation time to 0.85 s (the empirical mean pre-block fixation interval). We applied two adjustments, estimate + fixation time and estimate / (actual − fixation time), and re-ran the repeated-measures ANOVA (Supplement S3).

Session-position analyses were run in two parts (Supplement S4). For Watching and Baseline, we ran a 2 (Condition) × 2 (Position: second, third) repeated-measures ANOVA on participant means of the raw estimate. Two checks followed: the same ANOVA on the estimation ratio, and block-level linear mixed models. The paired Scrolling-Watching difference was computed within each repetition, and the Condition × Repetition interaction was tested by a likelihood-ratio test of block-level mixed models (participant random intercept, block length as covariate). The difference was also recomputed without the first and the last condition sequence of the experiment to exclude the primacy and recent effects. We further modeled sequential assimilation to account for the context effects. It regressed each block’s estimation ratio on the preceding block’s ratio (weight *g*) with condition, block length, and repetition as factors.

### ERP analysis

We computed picture-locked grand-average ERPs from the picture-onset epochs at a posterior-occipital cluster that joins the standard posterior (PO3, PO4, POz, PO7, PO8) and occipital (O1, Oz, O2) electrode groups over visual cortex, where picture-evoked components are maximal. The same cluster and windows were used for every picture-locked contrast. The natural-image waveform was dominated by a broad posterior positivity that did not form a clean canonical P1-N1-P2 sequence. We therefore used descriptive component labels: P1-like early positive (80–130 ms), P2-like posterior positive (140–300 ms), and the late positive potential (LPP, 300–500 ms). We take the P1-like window to index the initial sensory response of visual cortex to the picture, which is amplified when attention is directed to the stimulus (Hillyard & Anllo-Vento, 1998); the P2-like window to index the early selection and evaluation of picture content by the visual system (Schupp et al., 2006); and the LPP to index sustained attention to and elaborated processing of what the picture shows (Hajcak et al., 2010; Schupp et al., 2006). A difference between Scrolling and Watching in the early windows would therefore point to a change in how the picture was initially processed; a difference confined to the LPP would point to a change in how long it was elaborated. Mean amplitudes per participant, condition, and window served as the dependent variable for planned paired t-tests contrasting Scrolling and Watching.

The speed of the P2-like rise was measured as the 50%-rise latency: the first interpolated time point at which the waveform reached half its peak amplitude, with the peak taken in the 180-300 ms window. As a robustness check on the window means, a cluster-based permutation t-test (5000 permutations, α = .05, spatial adjacency from the 10–20 montage) covered the posterior cluster from 0 to 1000 ms. The full picture-locked analysis grid is in Supplement S5.

### FRP analysis

We computed fixation-related potentials (FRPs) from fixation-locked epochs, averaging them separately for Scrolling, Watching, and Baseline at the occipital cluster (O1, Oz, O2) used in previous FRP work on natural scenes (Madison et al., 2025). We focused on the lambda response, the early occipital visual evoked potential elicited when a saccade terminates and the retinal image refreshes, measuring mean amplitude in the 60–120 ms post-fixation window. Its amplitude indexes the visual input taken in at each fixation and scales with the amplitude of the preceding saccade.

As a stimulus-sampling check, we estimated the condition effect in a linear mixed model of single-fixation lambda amplitudes with crossed random intercepts for participant and image (statsmodels MixedLM, maximum likelihood). Because it has no by-condition random slopes, the model checks the condition mean rather than condition-by-item variation. Beyond lambda, three later fixation-locked windows were defined for exploratory analysis: N1FRP (120–200 ms), P2FRP (200–350 ms), and a late window (350–600 ms). These later-window contrasts are exploratory and uncorrected, and the full cluster-by-window grids are reported in Supplement S6.

## Results

### Subjective duration

Self-paced Scrolling and yoked Watching, the two sequential-picture conditions, both compressed subjective duration relative to the static-photograph Baseline, and estimates were shortest under Scrolling. Participants consistently underestimated block duration across all conditions (Figure 2A; mean actual duration 60.70 s). Mean estimated durations were 28.29 s (*SD* = 16.27) for Scrolling, 30.66 s (*SD* = 16.97) for Watching, and 39.16 s (*SD* = 22.87) for Baseline.

**Figure 2.**
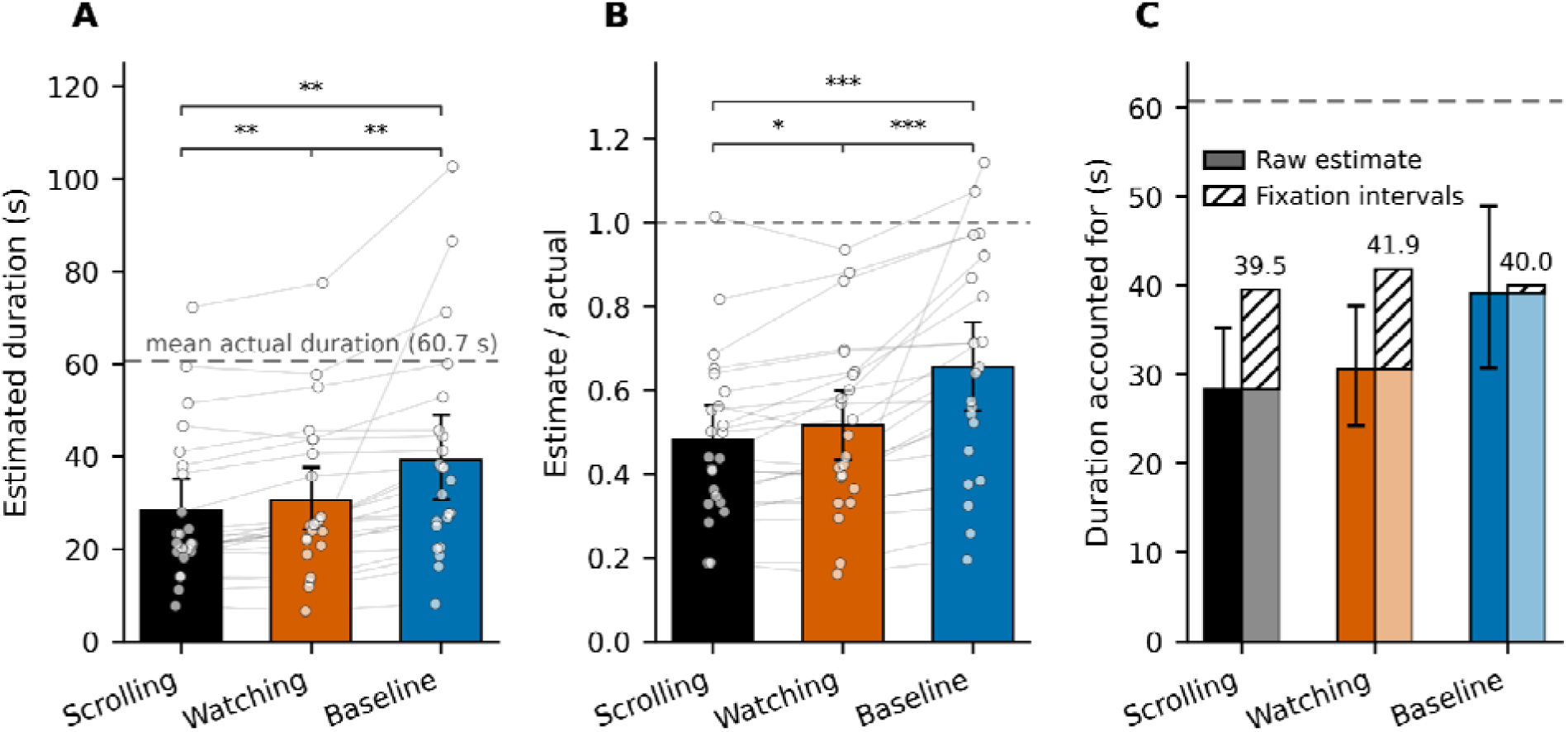
Subjective time compression and fixation intervals. Self-paced Scrolling compressed subjective duration relative to yoked Watching and to the static-photograph Baseline. (A) Raw estimated duration by condition (mean and 95% bootstrap CI, with paired subject lines). The dashed reference line marks the mean actual block duration (60.70 s). Scrolling was shorter than Watching (mean difference −2.37 s, *p*_Holm_ = .003), and both sequential-picture conditions were shorter than Baseline (Scrolling vs. Baseline −10.87 s, *p*_Holm_ = .003; Watching vs. Baseline −8.50 s, *p*_Holm_ = .005). (B) Estimation ratio (estimate divided by actual duration), anchored at 1.0 (veridical), reproducing the same ordering. (C) Raw estimate (filled bar) stacked with within-block fixation time (hatched segment). Once the inter-image fixation intervals are credited at their veridical weight, the condition effect on total accounted duration is no longer reliable, *F*(1.08, 23.72) = 0.58, *p* = .465, = .026. The accounted-for totals (roughly 40 s) remain well short of the 60.70 s actual duration (dashed line), consistent with participants underweighting the inter-image fixation intervals while still compressing the block as a whole. *N* = 23.

A 3 × 3 repeated-measures ANOVA (condition × block length) on raw estimates revealed a main effect of condition, *F*(1.08, 23.72) = 12.20, *p* = .002, 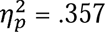 (Greenhouse–Geisser corrected), and block length, *F*(1.22, 26.85) = 42.97, *p* < .001, 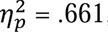, with no interaction, *F*(4, 88) = 1.25, *p* = .295. Holm-corrected paired contrasts confirmed all three predicted orderings: Scrolling vs. Watching (mean difference −2.37 s, 95% CI [−3.70, −1.04]; *t*(22) = −3.69, *p*_Holm_ = .003; *d*_z_ = −0.77, 95% CI [−1.23, −0.30]), Scrolling vs. Baseline (−10.87 s [−16.77, −4.97]; *t*(22) = −3.82, *p*_Holm_ = .003; *d*_z_ = −0.80 [−1.26, −0.32]), and Watching vs. Baseline (−8.50 s [−14.21, −2.80]; *t*(22) = −3.09, *p*_Holm_ = .005; *d*_z_ = −0.64 [−1.09, −0.19]). The between-participant *SD*s above reflect individual differences in estimation scale, not the precision of these paired contrasts: the within-participant correlation between Scrolling and Watching means was *r* = .98, so the *SD* of the paired Scrolling-Watching differences was 3.1 s. The baseline contrasts were less tightly paired but still precise (*r* = .81 and .82; *SD* of the differences 13.6 and 13.2 s).

Normalization via the estimation ratio (estimate / actual duration) mirrored these raw patterns (Figure 2B). Mean ratios were 0.481 (*SD* = 0.197) for Scrolling, 0.515 (*SD* = 0.210) for Watching, and 0.654 (*SD* = 0.260) for Baseline. A parallel ANOVA on the estimation ratio yielded a robust main effect of condition, *F*(1.20, 26.45) = 18.79, *p* < .001, = .461, and block length, *F*(1.74, 38.31) = 9.05, *p* = .001, = .292, with no interaction (*F* < 1). Holm-corrected pairwise contrasts again confirmed the same ordering, with ratios lowest under Scrolling, intermediate under Watching, and highest under Baseline (*t*s(22) < −2.53, *p*_Holm_ ≤ .019, *d*_z_ ≤ −0.53).

### Fixation intervals

Because an inter-image fixation interval preceded every picture, the sequential-picture blocks contained many short periods without picture content. We added these fixation intervals (11.21 s for Scrolling and Watching; 0.85 s for the modeled Baseline) to raw estimates to assess if duration judgments tracked the presence of visual content alone. This adjustment yielded mean accounted durations of 39.50 s for Scrolling, 41.87 s for Watching, and 40.01 s for Baseline (Figure 2C), effectively eliminating the condition effect, *F*(1.08, 23.72) = 0.58, *p* = .465, 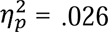. The adjustment left the Scrolling-Watching difference unchanged by design: the two conditions contained identical fixation-interval time, so whatever separates Scrolling from Watching lies outside the fixation intervals. A robustness check on ratio measures returned the same pattern. Removing fixation time from the actual duration before computing the ratio yielded fixation-removed ratios of 0.628 for Scrolling, 0.669 for Watching, and 0.666 for Baseline, with no reliable condition effect, *F*(1.31, 28.90) = 0.89, *p* = .380, 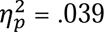 (Supplementary S3, Figure S3.1).

The fixation-interval adjustment did not restore veridical timing: even with these intervals fully credited, accounted-for durations fell roughly 20 s short of the 60.70 s actual block duration (Figure 2C).

### Session position and sequential context

Because Scrolling always came first with a repetition, and Watching and Baseline alternated between the second and third positions, position was testable directly for Watching and Baseline and only indirectly for Scrolling.

For Watching and Baseline, a 2 (Condition) × 2 (Position: second, third) repeated-measures ANOVA on participant means of the raw estimate (*N* = 22 with all four cells) gave a main effect of condition, *F*(1, 21) = 8.92, *p* = .007, 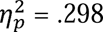, a main effect of position, *F*(1, 21) = 5.37, *p* = .031, 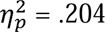 (third minus second: +2.1 s, 95% CI [0.2, 4.0]), and no reliable Condition × Position interaction, *F*(1, 21) = 1.92, *p* = .180, 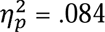. The shift was numerically carried by Watching (+5.5 s [−0.2, 11.2], *p*_Holm_ = .120) and not by Baseline (−1.3 s [−6.4, 3.8], *p*_Holm_ = .602). The estimation ratio and the block-level mixed models yielded similar results (interaction *p* = .010 and *p* = .043; Supplement S4, Table S4.1 and Figure S4.1A). Because the second position always followed Scrolling and the third always followed the other condition, this shift is as much an effect of the preceding condition as of ordinal position, and it does not bear on Scrolling’s own first position.

Scrolling’s first position was examined across repetitions (Supplement S4, Figure S4.2A). Scrolling came first in the first repetition as in every other, yet there Scrolling and Watching did not differ (−0.30 s [−1.96, 1.35], *d_z_* = −0.08). The difference appeared from the second repetition on (−5.23 s [−9.89, −0.57], −1.77 s [−5.41, 1.88], and −2.62 s [−5.46, 0.21] in repetitions 2–4), although the Condition × Repetition interaction was not reliable, χ^2^ (3) = 2.01, *p* = .570. An effect of being first as such does not predict this pattern, since it would act in the first repetition too. The Scrolling − Baseline difference was present in every repetition (9.9 to 12.2 s, all *p* ≤ .006). Overall estimates rose from the first to the second repetition (+9.8 s [2.4, 17.2]) and did not rise further.

Together, these results point to a sequential effect: each estimate was drawn toward the one before it. Watching was preceded either by Scrolling, judged short, or by Baseline, judged long, whereas Baseline was preceded by two conditions judged alike, so only Watching should move with position. We fitted a post hoc block-level model in which each block’s estimation ratio is pulled toward the preceding block’s ratio. It gave a weight of *g* = 0.24 [0.17, 0.31] and reproduced this asymmetry in direction, accounting for about half of the Watching shift (Supplement S4, Figure S4.1B). The same pull shapes the self-paced contrast. The Scrolling − Watching difference was absent when Watching immediately followed Scrolling (−0.17 s [−1.87, 1.53]) and about 5 s when Watching followed Baseline (−4.79 s [−7.07, −2.51]; Figure S4.2A): next to Scrolling, Watching is drawn toward Scrolling’s short estimate, and after Baseline toward Baseline’s long one. This pattern is consistent with a shortening under self-paced advance that assimilation masks in one context and enlarges in the other. These analyses, however, cannot fully rule out a contribution of Scrolling’s fixed first position, which from the second repetition on also followed a break and eye-tracker recalibration. We return to this in the Limitations.

### Eye movements

Eye movements documented the behavioral context of self-paced viewing (Figure 3). Mean fixation duration was shortest under Scrolling (307.6 ms, *SD* = 53.8), intermediate under Watching (318.5 ms, *SD* = 70.8), and longest under Baseline (354.1 ms, *SD* = 95.5). A repeated-measures ANOVA yielded a reliable condition main effect, *F*(1.38, 30.41) = 8.00, *p* = .004, 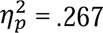 (Greenhouse-Geisser corrected). Both sequential-picture conditions produced shorter fixations than Baseline (Scrolling vs. Baseline: *t*(22) = −2.96, *p*_Holm_ = .014, *d*_z_ = −0.62, 95% CI [−79.0, −13.9 ms]; Watching vs. Baseline: *t*(22) = −3.71, *p*_Holm_ = .004, *d*_z_ = −0.77, 95% CI [−55.4, − 15.7 ms]), but Scrolling and Watching did not differ reliably (*t*(22) = −1.07, *p*_Holm_ = .298, *d*_z_ = −0.22). Sequential-picture viewing therefore shortened individual fixations relative to the static photograph, yet self-paced control shortened them no further.

**Figure 3.**
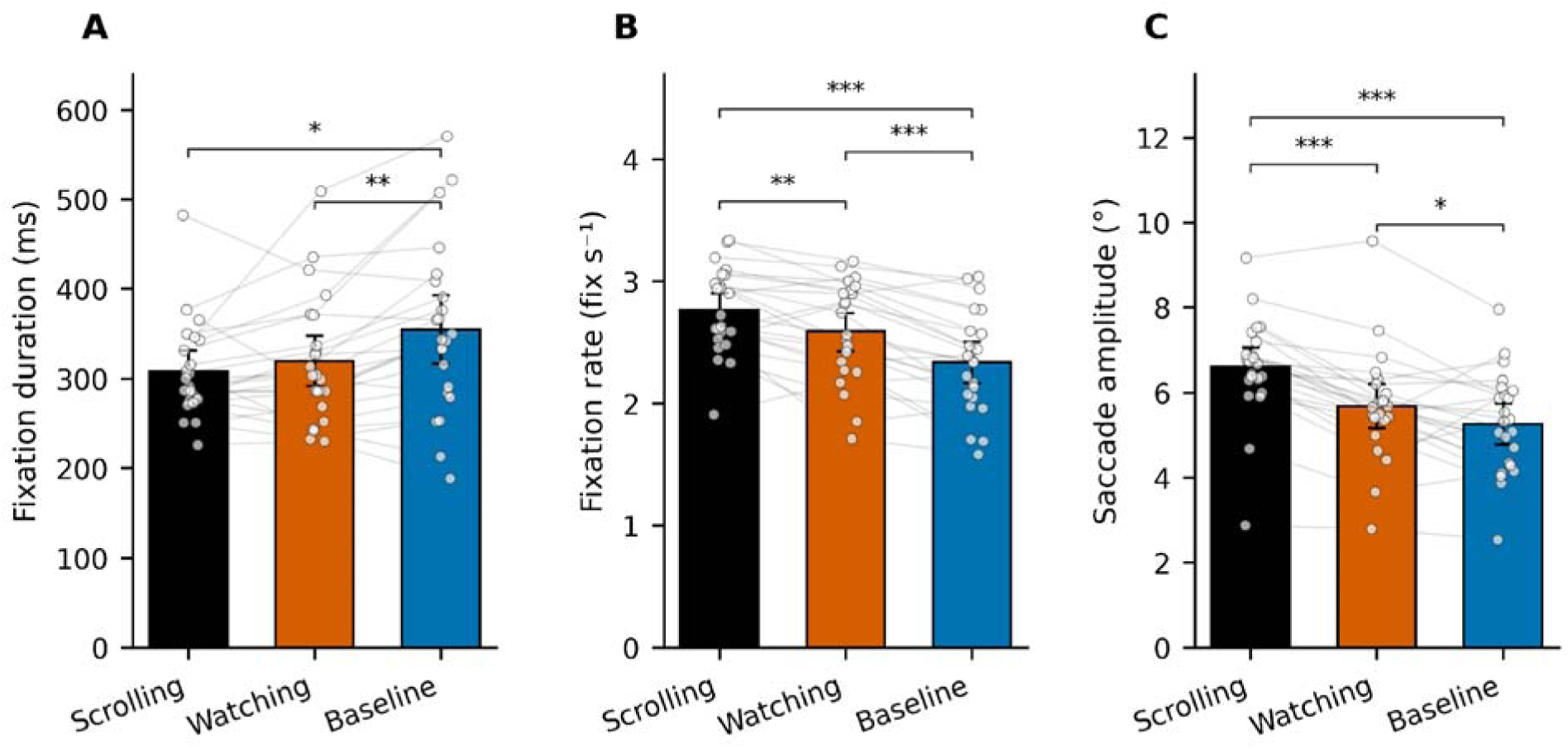
Oculomotor measures by condition. (A) Mean fixation duration was shorter under sequential-picture viewing than under the static-photograph Baseline, but did not differ reliably between Scrolling and Watching (*d*_z_ = −0.22, *p*_Holm_ = .298). The Scrolling-Watching time difference therefore did not depend on shorter individual fixations. (B) Fixation rate showed the full active-control gradient (Scrolling > Watching > Baseline; all *p*_Holm_ ≤ .003). (C) Mean saccade amplitude in degrees of visual angle followed the same gradient, with Scrolling producing the broadest spatial sampling (6.60° vs. 5.68° vs. 5.26°; all contrasts *p*_Holm_ ≤ .042). Bars show condition means with 95% bootstrap CIs; lines and points show individual subjects. Stars mark Holm-corrected contrasts (* *p* < .05, ** *p* < .01, *** *p* < .001). *N* = 23.

Fixation rate followed the reverse ordering: 2.76 fix/s (*SD* = 0.35) under Scrolling, 2.59 fix/s (*SD* = 0.41) under Watching, and 2.33 fix/s (*SD* = 0.41) under Baseline. The condition main effect was large, *F*(2, 44) = 29.04, *p* < .001, 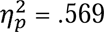, and all three pairwise contrasts were reliable (Scrolling vs. Watching: *d*_z_ = 0.70; Scrolling vs. Baseline: *d*_z_ = 1.30; Watching vs. Baseline: *d*_z_ = 1.15; all *p*_Holm_ ≤ .003). Self-paced control produced the highest fixation rate, and yoked passive viewing remained above the static-photograph Baseline.

Mean saccade amplitudes followed the same gradient as the rate measures: Scrolling 6.60° (*SD* = 1.19), Watching 5.68° (*SD* = 1.29), Baseline 5.26° (*SD* = 1.19). The condition main effect was highly significant, *F*(2, 44) = 32.43, *p* < .001, 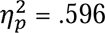, and all three contrasts were reliable (Scrolling vs. Watching: *t*(22) = 7.29, *p*_Holm_ < .001, *d*_z_ = 1.52; Scrolling vs. Baseline: *t*(22) = 7.28, *p*_Holm_ < .001, *d*_z_ = 1.52; Watching vs. Baseline: *t*(22) = 2.16, *p*_Holm_ = .042, *d*_z_ = 0.45). Self-paced viewers made larger saccades per picture advance than participants under yoked or static-photograph viewing, consistent with broader spatial sampling of each scene.

### Picture-locked ERP

Across the posterior-occipital cluster, baseline-corrected amplitudes were numerically larger under Scrolling than under Watching in all three windows (Figure 4A). The P1-like early positive was small but cleared the threshold for the planned Scrolling–Watching contrast (3.08 µV vs. 2.67 µV), *t*(22) = 2.10, *p* = .048, *d*_z_ = 0.44, 95% CI [0.004, 0.83]. The P2-like posterior positive carried the clearest difference (9.76 µV vs. 8.84 µV), *t*(22) = 4.11, *p* < .001, *d*_z_ = 0.86, 95% CI [0.45, 1.37], and the LPP window followed (7.63 µV vs. 6.90 µV), *t*(22) = 2.84, *p* = .010, *d*_z_ = 0.59, 95% CI [0.20, 1.27].

**Figure 4.**
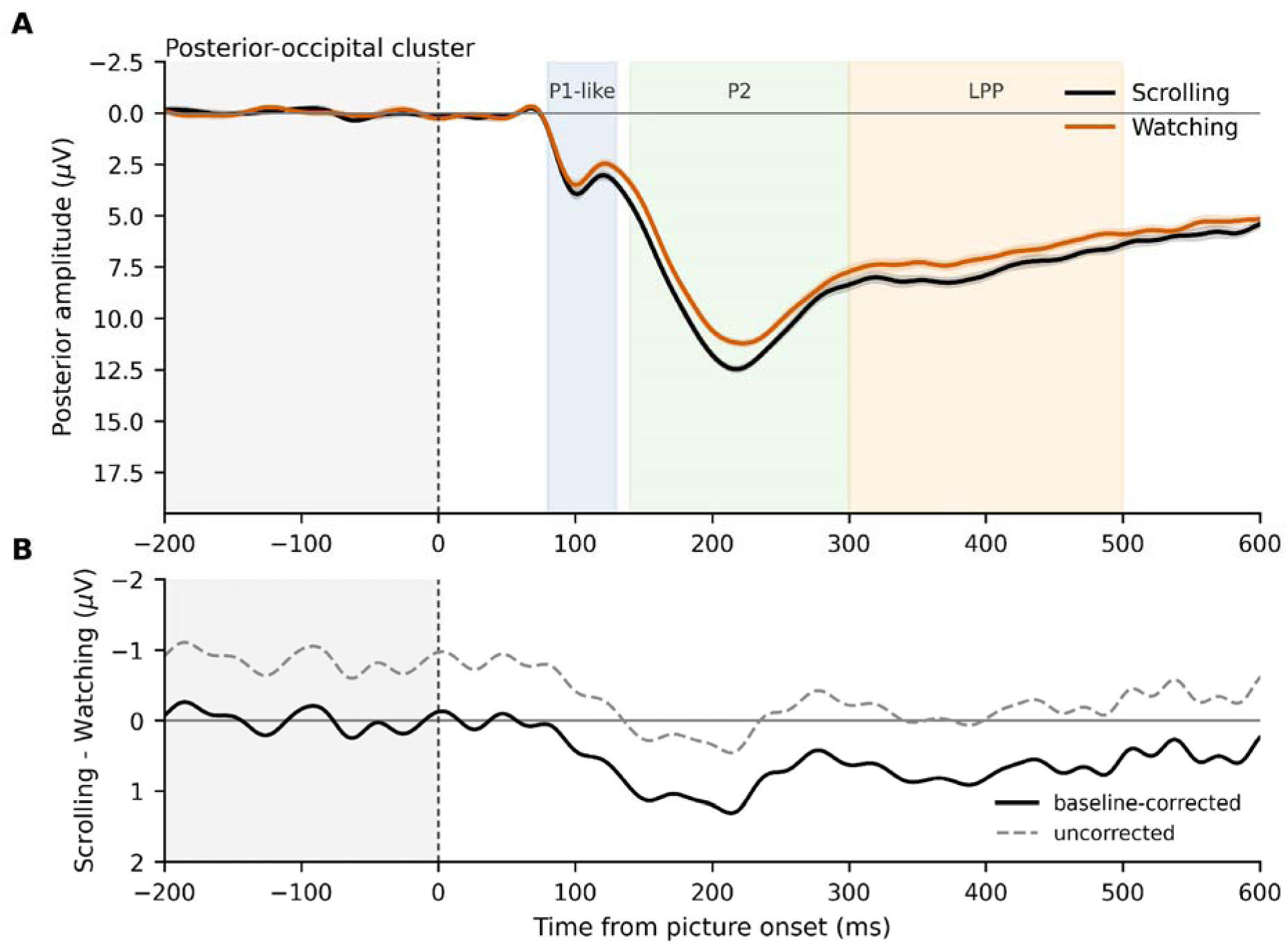
Picture-locked posterior ERP. (A) Baseline-corrected grand-average ERP at the posterior-occipital cluster (PO3, PO4, POz, PO7, PO8, O1, Oz, O2) from −200 to 600 ms relative to picture onset, with Scrolling (black) and Watching (red-orange) traces and within-subject 95% CI ribbons. The gray band marks the pre-onset baseline window; coloured bands mark the P1-like early positive (80–130 ms), P2-like posterior positive (140–300 ms), and LPP (300–500 ms) windows used for amplitude extraction. (B) Scrolling-minus-Watching difference wave from the same posterior-occipital cluster. The solid trace shows the baseline-corrected difference; the dashed trace shows the uncorrected difference after restoring the pre-onset Scrolling-minus-Watching offset. Baseline-corrected amplitudes were larger under Scrolling in the P1-like early positive, P2-like posterior positive, and LPP windows, but the corresponding uncorrected post-onset differences were not reliable. The rise toward the P2-like posterior positivity reached its 50%-rise point earlier under Scrolling than Watching (135.3 vs. 141.1 ms), *t*(22) = −3.02, *p* = .006, *d*_z_ = −0.63. *N* = 23.

The breadth of the baseline-corrected difference raised a question about the baseline itself, since picture onsets were self-triggered by mouse click under Scrolling and externally imposed under Watching. We therefore re-epoched the same retained picture events from the cleaned continuous recordings without pre-stimulus baseline correction. In the uncorrected −200 to 0 ms pre-onset window, posterior-occipital amplitude was more negative under Scrolling than under Watching (−3.46 µV vs. −2.61 µV), *t*(22) = −2.21, *p* = .038, *d*_z_ = −0.46, whereas over central scalp sites near the motor electrodes the same pre-onset contrast was not reliable, *t*(22) = −0.90, *p* = .380.

Without baseline correction, the Scrolling-versus-Watching differences were not reliable for the P1-like early positive (−0.43 µV, *p* = .274), P2-like posterior positive (+0.05 µV, *p* = .906), or LPP (−0.14 µV, *p* = .743). Because baseline correction subtracts each condition’s pre-onset mean, the more negative Scrolling baseline carries forward as a positive Scrolling-minus-Watching offset after onset. The corrected between-condition difference therefore indexed an anticipatory or carry-over posterior state present before picture onset, not enhanced encoding of the picture under self-paced control.

A rise-latency analysis added a separate timing measure. The P2-like posterior positivity reached its 50%-rise point earlier under Scrolling than under Watching (135.3 ms vs. 141.1 ms), *t*(22) = −3.02, *p* = .006, *d*_z_ = −0.63, 95% CI [−0.90, −0.32]. This result converges on the same reading: self-paced onset shifts the timing of the early posterior response rather than amplifying the picture response itself. Full component ERP contrasts, rise-latency values, and the cluster-permutation diagnostic appear in Supplement S5.

### Fixation-locked FRP

Fixation-locked occipital responses followed the active-control manipulation in graded fashion, with the lambda response strongest under Scrolling, intermediate under Watching, and smallest under Baseline (Figure 5). Mean lambda amplitudes in the occipital cluster (O1, Oz, O2) were 3.71 µV under Scrolling, 3.44 µV under Watching, and 2.84 µV under Baseline. A one-way repeated-measures ANOVA returned a reliable condition effect, *F*(1.49, 32.79) = 22.89 *p* < .001, = .510 (Greenhouse-Geisser corrected). Holm-corrected pairwise contrasts then ordered the conditions cleanly: Scrolling exceeded both Watching and Baseline, and Watching exceeded Baseline (*t*s > 3.10, *p*_Holm_ < .004, *d*_z_ > 0.66).

**Figure 5.**
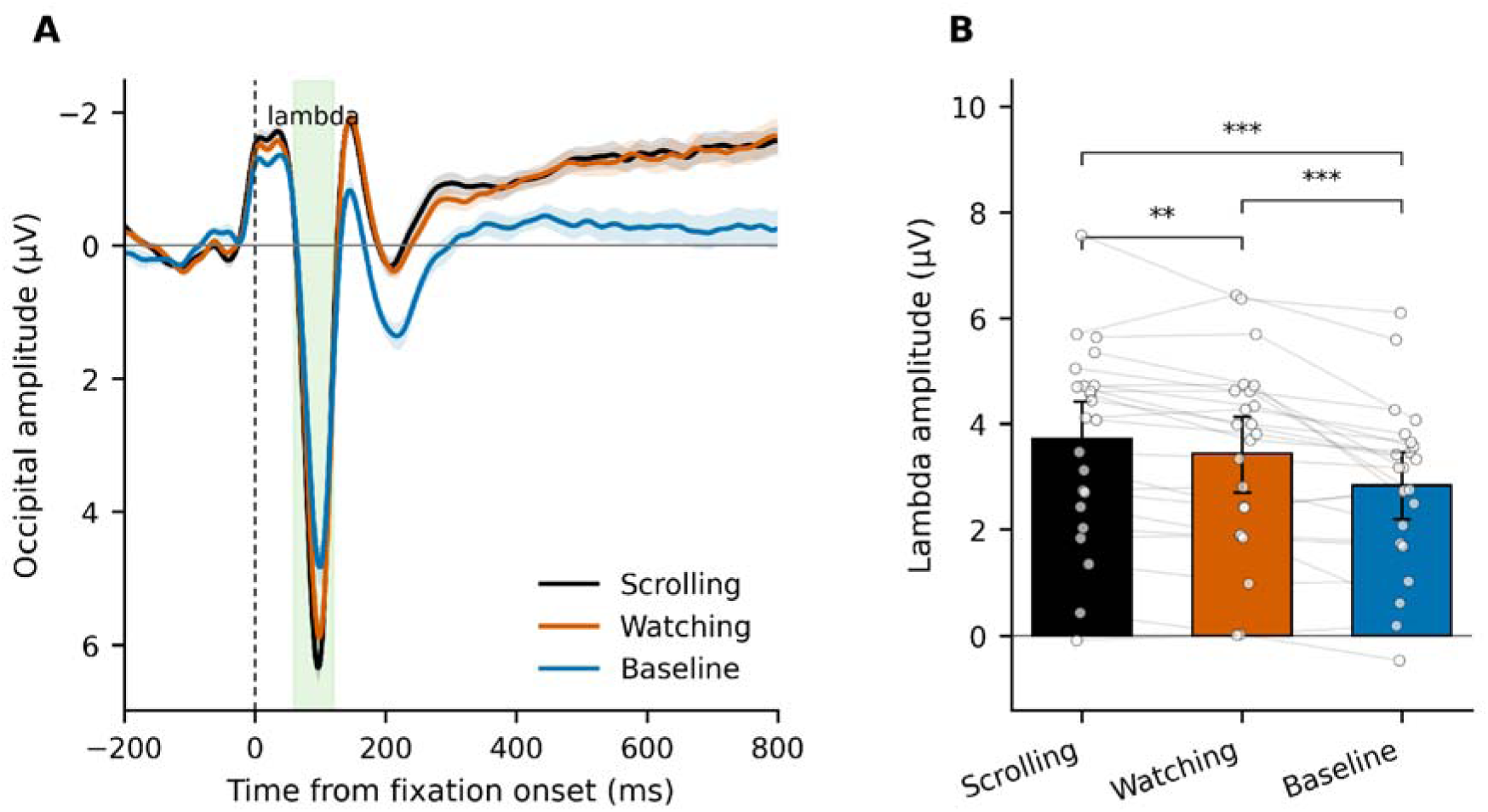
Fixation-locked occipital FRP. The occipital lambda was strongest under Scrolling, intermediate under Watching, and smallest under Baseline. (A) Grand-average fixation-locked potentials at the occipital cluster (O1, Oz, O2) under Scrolling (black), Watching (red-orange), and Baseline (blue), with within-subject 95% CI ribbons and the lambda window (60–120 ms) shaded. (B) Subject-mean lambda amplitude by condition, with within-subject paired lines, individual subject points, and a condition mean with 95% bootstrap CI. Mean lambda amplitudes were 3.71 µV (Scrolling), 3.44 µV (Watching), and 2.84 µV (Baseline), differing in the order, with Scrolling above Watching (*t*(22) = 3.19, *p*_Holm_ = .004, *d*_z_ = 0.66), Scrolling above Baseline (*t*(22) = 5.62, *p*_Holm_ < .001, *d*_z_ = 1.17), and Watching above Baseline (*t*(22) = 4.16, *p*_Holm_ < .001, *d*_z_ = 0.87). FRPs were recomputed after an eye-tracker-to-EEG synchronization correction. *N* = 23.

The Scrolling-versus-Watching lambda contrast also survived a single-fixation mixed model that treated participants and images as crossed random samples, *b* = 0.227 µV, SE = 0.060, *t* = 3.80, *p* < .001. The same model showed no comparable Scrolling-versus-Watching enhancement in the later windows: the contrast was null at N1_FRP_ (*p* = .349) and the late window (*p* = .745), and P2_FRP_ carried only a small difference in the opposite direction (*b* = −0.14 µV, *p* = .039). Both sequential-picture conditions nonetheless diverged sharply from Baseline across N1_FRP_, P2_FRP_, and the late window (all *p* < .001), so these windows registered robust fixation-evoked activity that did not track self-paced control (Supplement S6). The effect of active control is thus confined to an early fixation-locked lambda modulation rather than a broad amplification of all fixation-locked activity under self-paced viewing.

The lambda is a saccade-evoked transient whose amplitude scales with the size of the retinal-image shift at each fixation. Self-paced viewing produced both more saccades and larger ones, and the larger early lambda per fixation tracked these larger saccades. Adding incoming saccade amplitude to the single-fixation model reduced the Scrolling-versus-Watching lambda coefficient from 0.20 to 0.05 µV and rendered it non-significant, *p* = .40, while saccade amplitude was itself a strong predictor at 0.23 µV per degree (Supplement S6). The saccade-amplitude covariate thus statistically accounted for the Scrolling-versus-Watching lambda difference, as expected if the difference reflects how the eyes sampled the scene (more and larger saccades onto fresh content). A covariate adjustment of this kind does not establish that saccade amplitude mediated the condition effect.

### Recognition

Mean recognition accuracy was highest under Baseline (0.996, *SD* = 0.017), followed by Scrolling (0.913, *SD* = 0.089) and Watching (0.880, *SD* = 0.117). Both sequential-picture conditions fell well below Baseline (*t*s < −4.74, *p*s < .001). A Baseline block presented a single static photograph, so we treated Baseline as a sanity check on the probe and constrained the informative comparison to the two sequential-picture conditions. A 2 (condition: Scrolling, Watching) × 3 (block length: 9, 12, 18 pictures) repeated-measures ANOVA on accuracy returned no condition effect, *F*(1, 22) = 2.31, *p* = .142, 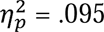 (the equivalent paired contrast gives *d*_z_ = 0.32, 95% CI [−0.11, 0.73]), with Scrolling only numerically above Watching (0.913 vs 0.880). Accuracy did decline with block length, *F*(2, 44) = 4.14, *p* = .022, 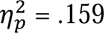, from 0.935 at nine pictures to 0.842 at eighteen, and the condition by block-length interaction was negligible, *F*(2, 44) = 0.11, *p* = .897, 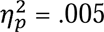. A two-one-sided-tests (TOST) equivalence test did not establish that Scrolling and Watching were equivalent within a smallest effect size of interest of *d*_z_ = ±0.50, *p* = .195. Because the probe was sparse (about 12 judgments per condition) and the equivalence test inconclusive, the null neither rules out a small difference nor establishes its absence.

Exploratory brain–behavior correlations between picture-locked posterior ERP amplitudes and the duration estimate did not survive false-discovery-rate correction.

## Discussion

Sequential-picture viewing compressed prospective time. Relative to a single static photograph shown for the same duration, blocks of self-paced Scrolling and of yoked Watching were judged shorter by roughly 8.5–11 s of a 60-s block (Figure 2). A descriptive decomposition locates most of this compression in the inter-image fixation intervals. Self-paced advance added a smaller compression: Scrolling was judged shorter than Watching in the paired analysis, but by an amount that depended on sequential context. The EEG and eye-tracking measures characterize what self-paced viewing did to concurrent processing: it changed the state in which each picture arrived and the way the eyes sampled it, without strengthening the picture-evoked response.

### Timing mechanism: attention to time and to information-bearing events

There are two accounts that make opposite predictions for a picture stream: an attentional gate, on which duration shrinks as attention leaves the clock (Polti et al., 2018; Zakay & Block, 1997; Zang et al., 2026), and a content readout, on which duration grows with the salient changes an interval contains (Roseboom et al., 2019; Sherman et al., 2022). Each is well supported in the setting its evidence comes from. The content readout has been demonstrated with continuous natural-scene videos and with filled versus empty intervals, where the salient changes themselves vary and are registered as the interval unfolds (Horr & Di Luca, 2015; Roseboom et al., 2019; Sherman et al., 2022). The present task asked for a minute-long prospective estimate while a stream of discrete pictures, and in Scrolling the decision when to advance, competed with time for attention. In this setting the results support the attentional account: sequential pictures felt shorter than a static photograph, not longer.

The fixation-interval analysis pointed to that potential source. Adding the fixation time back to the raw estimates brought Scrolling, Watching, and Baseline within roughly two seconds of one another (Figure 2C). Most of the Baseline difference therefore lay in the intervals punctuating the sequential streams rather than in the pictures themselves: observers timed the pictures and barely counted the intervals between them. On its own, then, the sequential-versus-Baseline compression likely comes from neglecting the fixation gaps, not from the pictures. Because the fixation interval was never manipulated independently of condition, this decomposition is arithmetic rather than a causal test, and even the credited estimates still fell some twenty seconds short of the 60.7 s objective duration.

Sequential viewing was nonetheless more active in its visual processing: Scrolling and Watching drew more and larger saccades than Baseline, with correspondingly larger eye-movement-locked occipital responses, yet this vigor did not feed the duration estimate of the pictures once the fixation intervals were excluded. The intensity of perceptual processing was dissociable from felt duration, and early evidence already suggested that the amount of information processed need not change duration judgments (Hicks et al., 1976).

The self-paced compression is suggested rather than established. Its direction held throughout, and because the Scrolling and Watching conditions carried identical fixation time, the difference does not depend on interval crediting. Its size, however, depended on sequential context in a pattern of assimilation, each estimate drawn toward the preceding one. That pattern fits a shortening under self-paced advance that assimilation masks when Watching follows Scrolling and enlarges when it follows Baseline. Two further contributors remain candidates.

One is onset predictability and self-generation: a self-triggered picture is anticipated, and at sub-second scales predictable stimuli are judged shorter while unexpected onsets expand judged duration (Kanai & Watanabe, 2006; Pariyadath & Eagleman, 2007), though the extension to minute-long estimates is untested. The other is the Scrolling sequence’s fixed place at the start of each repetition, directly after a break, which the design cannot separate from self-paced control (see Limitations and Outlook). An attention-weighted account accommodates a genuine control effect, since deciding when to advance, programming the saccade, selecting the dwell, and integrating the new picture draw on the same supervisory resources as monitoring elapsed seconds

Neither account in its pure form fits the whole pattern. A pure content readout predicts that the sequential-picture conditions should feel longer than the single photograph, yet they felt shorter; pure gating (Zakay & Block, 1997) would let attention return to the clock during the fixation intervals, yet those intervals were neglected. An attention-weighted readout reconciles both: duration reflects the interval’s content weighted by how much attention reaches time, high when nothing competes (the static photograph, judged longest) and low when content, control, or anticipation of the next picture draw attention away (the sequential stream and its intervals alike). Recent work proposes this single content-weighted process for timing (de Jong et al., 2025), on which the split between prospective, attention-based and retrospective, memory-based timing becomes a difference of degree (Brown, 1985) rather than a categorical distinction (Block & Zakay, 1997). The recognition probe, sparse as it is, gave no sign that the two sequential conditions differed in what was encoded, which offers little support for memory-based reconstruction as an explanation of their difference, even if recall shaped the absolute level of a minute-long estimate. The literature has independent support: understimulated intervals are reproduced as longer (Bangert et al., 2019), time feels more sluggish during passive waiting than task engagement (Meteier et al., 2025), and what matters is how an interval’s structure directs attention rather than whether it is filled (Church et al., 2007; Miki & Santi, 2005; Santi et al., 2005). The pervasive underestimate, present even for the static photograph, likely reflects measurement: prospective timing chronically underestimates long intervals, and central-tendency bias pulls estimates toward the session mean (Glasauer & Shi, 2022; Jazayeri & Shadlen, 2010; Shi et al., 2013).

### Self-paced control and the visual response

The behavioral effects raise a second question: did self-paced viewing strengthen visual processing of each picture? The baseline-corrected ERP seemed to say so, with a larger posterior positivity under Scrolling in all three windows (Figure 4). But Scrolling already sat below

Watching before picture onset, and without baseline correction the post-onset differences disappeared, as expected when the pre-stimulus interval itself differs between conditions (Alday, 2019), and without baseline correction the post-onset differences disappeared. The apparent enhancement therefore belongs to the pre-onset state, not to the picture-evoked response.

The pre-onset offset is still informative. It appeared in the planned posterior-occipital cluster but not over the central motor sites, which makes a simple mouse-click explanation unlikely. The more plausible source is anticipatory or carry-over activity: Scrolling onsets were self-triggered, whereas Watching onsets were externally imposed. Slow anticipatory negativity precedes expected and self-timed events (Brunia & van Boxtel, 2001), and self-generated stimuli, predictable and selected by the participant, are anticipated through forward models that modulate the cortical response they evoke (SanMiguel et al., 2013). On this reading, self-paced advance changed the neural state in which the next picture arrived. The rise-latency result fits: the P2-like posterior positivity reached its 50%-rise point a few milliseconds earlier under Scrolling, a modest effect limited to the planned contrast but in the same direction. When observers control the moment of advance, the posterior response reaches criterion sooner, without a larger picture-evoked response.

The one fixation-locked measure to separate Scrolling from Watching told the same story. Occipital lambda was largest in Scrolling, but lambda scales with the size of each incoming saccade (Dimigen et al., 2011), and the larger saccades of self-paced viewing statistically accounted for the difference. No later fixation window showed an enhancement (Supplement S6). The gradient is best read as indexing how actively the eyes sampled the scene, and gives no sign of a stronger response to each picture.

These measures characterize self-paced viewing, but they do not establish that it caused the duration difference. The exploratory brain-behavior correlation did not survive correction, so the anticipatory state is a working account of what accompanies self-paced advance rather than a measured mediator of time compression. The bounded reading is that self-paced advance altered anticipatory state, response timing, and the fixation-locked visual transient: the manner of exploration rather than the strength of encoding.

### Implications for human-technology interaction

The laboratory analogue targets one affordance of scrolling, the self-paced visual advance through a stream of pictures, and turns a familiar complaint about time disappearing in feeds into a measurable effect. Self-report studies of social-media use describe absorption, reduced time awareness, and patchy memory for consumed content (Baughan et al., 2022; Sharma et al., 2022), trace-data work shows that adolescents misestimate their own smartphone use across weeks (Marciano & Camerini, 2022), and short-form video research has begun to measure in-task time distortion directly (Jiang et al., 2025; Yang et al., 2024), with effects whose direction depends on whether estimates are prospective or retrospective. Meteier et al. (2025) showed that engagement, rather than digitality on its own, separates active task time from passive waiting.

Stripped of reward, social signal, recommender algorithms, and personalized content, our task still finds a minute of scrolling-like viewing judged at about half its length, some ten seconds shorter than a minute spent with a single photograph. The relevance is to time awareness. People misjudge how long they have been looking; the seams between items barely register. This matters for a person’s ability to act on their own intentions about when to stop, whatever they choose to do with their time. The study shows that a perceptual affordance shared by every feed, before any platform-specific factor is added, moves the felt clock.

The inter-image interval was built to resemble the pauses that separate items in a feed, and observers barely counted it. We read this as evidence that attention is spread unevenly across a viewing interval: the pictures draw it, and the stretches between them, when nothing new is yet on screen, are credited little. Feeds are rarely seamless, even if their seams are shorter and less regular than ours. Content loads after a swipe, a clip pauses before it plays, transitions and inserted advertisements separate posts, and the viewer’s attention is already moving to the next item while these pass. We propose that such seams are discounted in the same way, and that across a session their uncounted time accumulates, so that the clock time a feed takes outruns the time a user feels has passed. This is one route to the misestimation of use seen in trace data (Marciano & Camerini, 2022). The proposal makes a prediction that our design could not test, because the interval was not varied: the compression should grow with the time the seams occupy and shrink where a stream has none. The direct test is to lengthen, shorten, or fill the interval, and to include gap-free viewing.

### Limitations and Outlook

Twenty-three participants is a small sample for a single experiment, and the confidence intervals say what it supports. It supports the within-participant ordering: the within-person correlation between Scrolling and Watching means was high, so the paired contrasts are precise, and the sequential-versus-Baseline differences are Holm-significant with confidence intervals that exclude zero in every sample. The Scrolling-versus-Baseline difference is also unchanged whichever position Baseline occupied. However, it does not resolve small secondary effects. The size of the self-paced effect is the clearest case. On average, Scrolling was judged 2.4 s shorter than Watching, with a 95% confidence interval from about 1 to 4 s. Once we considered the sequential effect, however, the difference depended on which condition Watching followed: it was essentially absent when Watching came directly after Scrolling and about 5 s when Watching came after Baseline. The sample therefore establishes the direction of the effect better than its actual size. The recognition probe, with about twelve judgments per condition, could neither detect nor exclude a moderate memory cost. We report precision through these intervals because a post hoc power calculation would not add information beyond them.

Order is a fixed feature of the design. Scrolling always came first within a repetition because the yoked durations must exist before they can be replayed. The order mattered through sequential assimilation, the likely reason Watching moved with position and Baseline did not (Figure S4.1). The Condition × Position interaction was reliable for the estimation ratio and in the block-level models but not reliable in the participant-level ANOVA on raw estimates, and the assimilation model reproduced the asymmetry in direction and about half in size. Sequential assimilation is therefore a supported but not an established explanation of the order effect.

Scrolling’s own position can only be examined across repetitions. Scrolling was first in the first repetition too, and there it did not differ from Watching. The difference appeared only from the second repetition on. This late onset argues against an effect of being first as such and against drift across the session, but it does not fully rule out a first-position contribution, because from the second repetition on every Scrolling sequence followed the break and eye-tracker recalibration that closed the preceding repetition.

Generality is bounded by who was tested: 23 young adults at a single European university, aged 22 to 36, characterized only by age and gender, whose habitual media use we did not measure, in a task that set aside the reward signals, social meaning, and algorithmic curation of real feeds. We expect the qualitative pattern wherever attention must be divided between looking and timing, but its magnitude across age, culture, and media habit remains open.

Three potential studies follow directly. Manipulating the fixation interval, including gap-free viewing, would turn the fixation-interval accounting from a decomposition into a test. A position-balanced or randomized order, with yoked durations taken from a separate calibration phase and with externally triggered but predictable onsets alongside self-triggered ones, would separate self-paced control from sequential context and from the predictability of onsets. And a direct replication would settle the size of the self-paced increment that this sample leaves between zero and about five seconds.

## Conclusion

Looking up from a feed to find that minutes have vanished is usually blamed on the platform, its rewards, its social pull, its algorithm. Our analogue stripped all of those away, and the time still went missing. It went missing most in sequential-picture viewing: a minute of pictures was judged some ten seconds shorter than a minute spent with one photograph, as if people timed the pictures and the intervals between them barely counted. Scrolling compressed time a little further than Watching, in a direction that held throughout, by an amount that depended on what had come just before. The thief is attention: looking, bracing for the next picture, and perhaps deciding when to move on all draw it away from the clock, and duration shrinks to whatever attention is left to count.

## Author contributions

C.Q. and A.Z. contributed equally and share first authorship. C.Q.: investigation, formal analysis, writing. A.Z.: methodology, investigation, formal analysis, writing. S.C.: methodology, investigation, writing. Z.S.: conceptualization, methodology, formal analysis, supervision, funding acquisition, writing. All authors were involved in writing and reviewing the manuscript. The authors used large-language-model assistants (Anthropic Claude and OpenAI Codex) to help draft and edit prose and analysis code and to check citations and statistics during preparation and revision. All study design, data collection, analysis decisions, interpretation, and final wording are the authors’ own, and the authors verified all AI-assisted output.

## Data and code availability

The behavioral data, the analysis scripts, and the per-figure regeneration scripts are openly available at https://github.com/msenselab/self-paced-viewing-compresses-time; the raw EEG and eye-tracking recordings are available from the corresponding author on request.

## Author Note

### Funding

This project was supported by Deutsche Forschungsgemeinschaft (DFG) project SH 166/10-1 to Zhuanghua Shi.

### Author Contributions

C.Q. and A.Z. contributed equally and share first authorship. C.Q.: investigation, formal analysis, writing. A.Z.: methodology, investigation, formal analysis, writing. S.C.: methodology, investigation, writing. Z.S.: conceptualization, methodology, formal analysis, supervision, funding acquisition, writing.

## Footnotes

1 one for technical failure (very high EEG impedance and eye-tracker failure), two whose Scrolling-Watching differences in raw estimates lay more than 2 SD from the mean of all participants, and one who misunderstood the task. Sensitivity checks in the Results retain the last participant retained (*N* = 24), or all three (*N* = 26).

