## Supplementary Materials for "The Attentional Thief: Scrolling-Like Picture Streams Compress Subjective Time"

This supplement gives the checks behind the main claims. Three within-participant conditions were compared: Scrolling (self-paced advance through photographs by mouse click), Watching (different photographs with the picture and fixation durations of the participant's own Scrolling block, jointly permuted), and a static-photograph Baseline (one photograph shown for the yoked block duration). In the two sequential-picture conditions each picture was preceded by an inter-image fixation interval (0.7–1.0 s). The sections follow the main text: image statistics (S1), retained EEG epochs (S2), fixation-interval accounting (S3), session position and exclusion sensitivity (S4), picture-locked ERPs (S5), and fixation-locked FRPs (S6). In every figure, points are paired differences (first minus second condition) or model coefficients, and error bars are 95% confidence intervals; *N* = 23 unless stated otherwise.

### S1. Low-level image statistics

The photographs actually presented (8,257 picture instances) did not differ between conditions on any of four whole-image statistics, and no statistic interacted with block length (Table S1.1). The check covers low-level features only. Semantic complexity was not rated.

Table S1.1. Whole-image statistics of the presented photographs by condition (participant-cell means averaged across block lengths), with the condition main effect and the Condition × Block-length interaction from a 3 × 3 repeated-measures ANOVA. *N* = 23.

| **Statistic** | **Scrolling** | **Watching** | **Baseline** | **F(2, 44)** | **p** | **Condition × Block length p** |
| --- | --- | --- | --- | --- | --- | --- |
| Mean luminance (0–1) | 0.224 | 0.226 | 0.234 | 1.31 | .280 | .701 |
| RMS contrast | 1.093 | 1.085 | 1.077 | 0.18 | .836 | .705 |
| Canny edge density | 0.098 | 0.099 | 0.102 | 0.85 | .434 | .738 |
| Spatial-frequency slope | −1.325 | −1.322 | −1.313 | 0.70 | .503 | .253 |

### S2. Retained EEG epochs

Of 7,437 picture-locked epochs, 6,674 (89.7%) entered the participant averages (Figure S2.1). Three participants lost more than 30% of their picture epochs (39.8%, 40.7%, 47.1%). After inspection their remaining counts supported condition averages, and they were retained. Baseline presents one picture per block, so its picture-locked average rests on at most 12 epochs per participant, which is why the picture-locked contrasts in S5 compare Scrolling and Watching only. Fixation-locked epochs were plentiful in all three conditions.

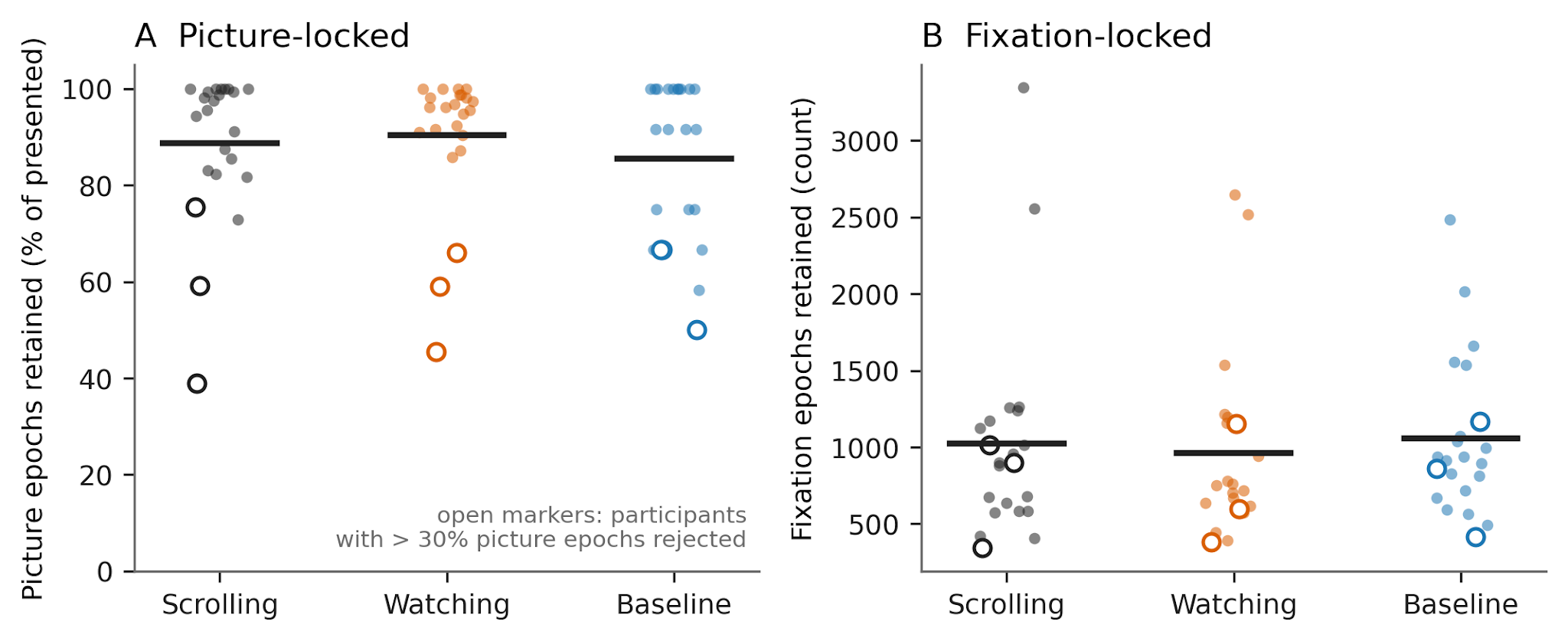

**Figure S2.1.** *Retained EEG epochs per participant.* (A) Percentage of presented picture epochs retained after AutoReject with a ±150 µV fallback. Mean retained counts were 140 (range 62–159) for Scrolling, 140 (69–156) for Watching, and 10 (6–12) for Baseline. (B) Retained fixation epochs after the fixation-duration filter and a ±150 µV criterion; means 1,024 (342–3,345), 963 (383–2,646), and 1,058 (418–2,484). Points are participants, bars are means, and open markers are the three participants with more than 30% of picture epochs rejected. Per-participant counts are in the project repository.

### S3. Fixation-interval accounting

Sequential-picture blocks contained a fixation interval before every picture (summed per block: 7.8, 10.3, and 15.5 s for 9, 12, and 18 pictures, identical in Scrolling and Watching), whereas Baseline had a single pre-block fixation (0.85 s). Crediting this time to the estimate, or removing it from the elapsed duration before computing the ratio, removed both Baseline contrasts and left the Scrolling − Watching contrast unchanged (Figure S3.1). The condition main effect was no longer reliable for either adjusted measure (estimate plus fixation intervals: *F*(1.08, 23.72) = 0.58, *p* = .465; fixation-removed ratio: *F*(1.31, 28.90) = 0.89, *p* = .380; Greenhouse–Geisser corrected), whereas block length still lengthened estimates.

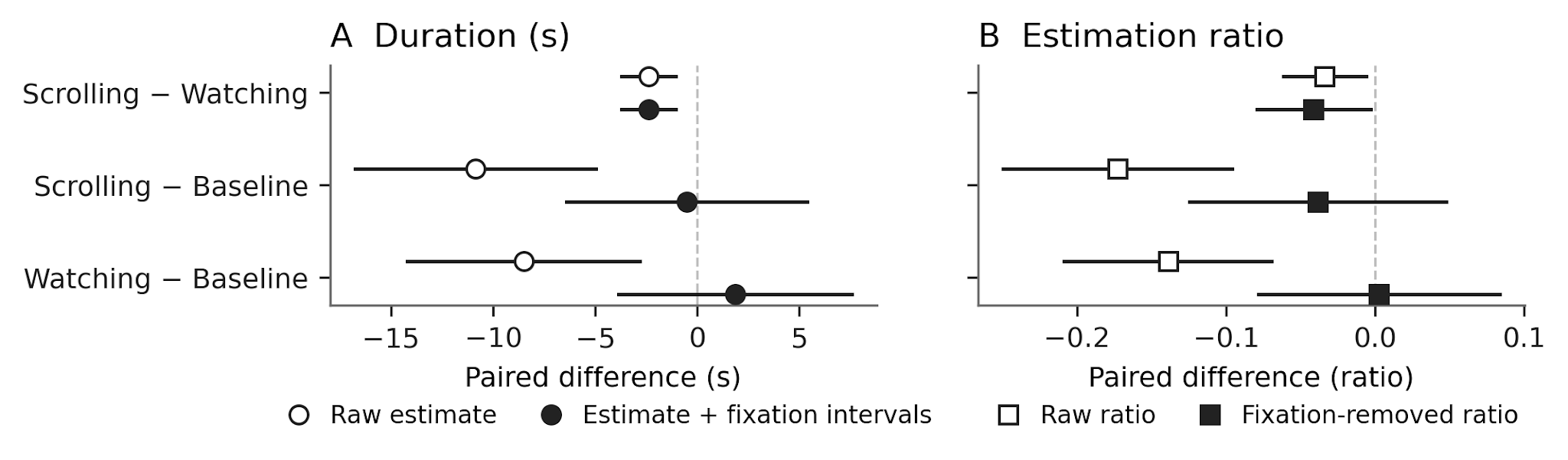

**Figure S3.1.** *Paired contrasts before and after fixation-interval accounting.* (A) Raw estimate (open) and estimate plus the block's summed fixation intervals (filled). (B) Raw estimation ratio (open) and the ratio computed against elapsed time minus the fixation intervals (filled). Scrolling and Watching had identical fixation-interval time, so their difference is the same before and after the adjustment.

### S4. Session position, sequential context, and exclusion sensitivity

Scrolling came first in every repetition because the Watching and Baseline blocks took their durations from it. Watching and Baseline alternated between the second and third positions (15 participants had Watching second in odd repetitions, 8 in even ones). The second position therefore always meant "immediately after Scrolling". The yoking held in the logged data: in all 267 Scrolling–Watching block pairs the picture and fixation durations were the same set, no photograph was shared, and total block durations matched within 0.6 s. One retained participant completed only the first repetition and is absent from analyses that need both positions (*n* = 22 there).

For Watching and Baseline, a 2 (Condition) × 2 (Position) repeated-measures ANOVA on the raw estimate gave main effects of condition and position and no reliable interaction (Table S4.1). The shift sat in Watching, which was judged longer when it came third, after Baseline, than second (Figure S4.1A). The shift was reliable for the ratio and in block-level mixed models that keep all blocks of all 23 participants (Condition × Position *b* = −6.3 s [−12.5, −0.2], *p* = .043), and the same ANOVA on actual block duration showed no position effect (*p* = .345), so the ratio result does not come from its denominator. As a consequence, the Scrolling − Watching difference was absent when Watching immediately followed Scrolling and about 5 s when Watching followed Baseline (Figure S4.2A), in both counterbalancing groups.

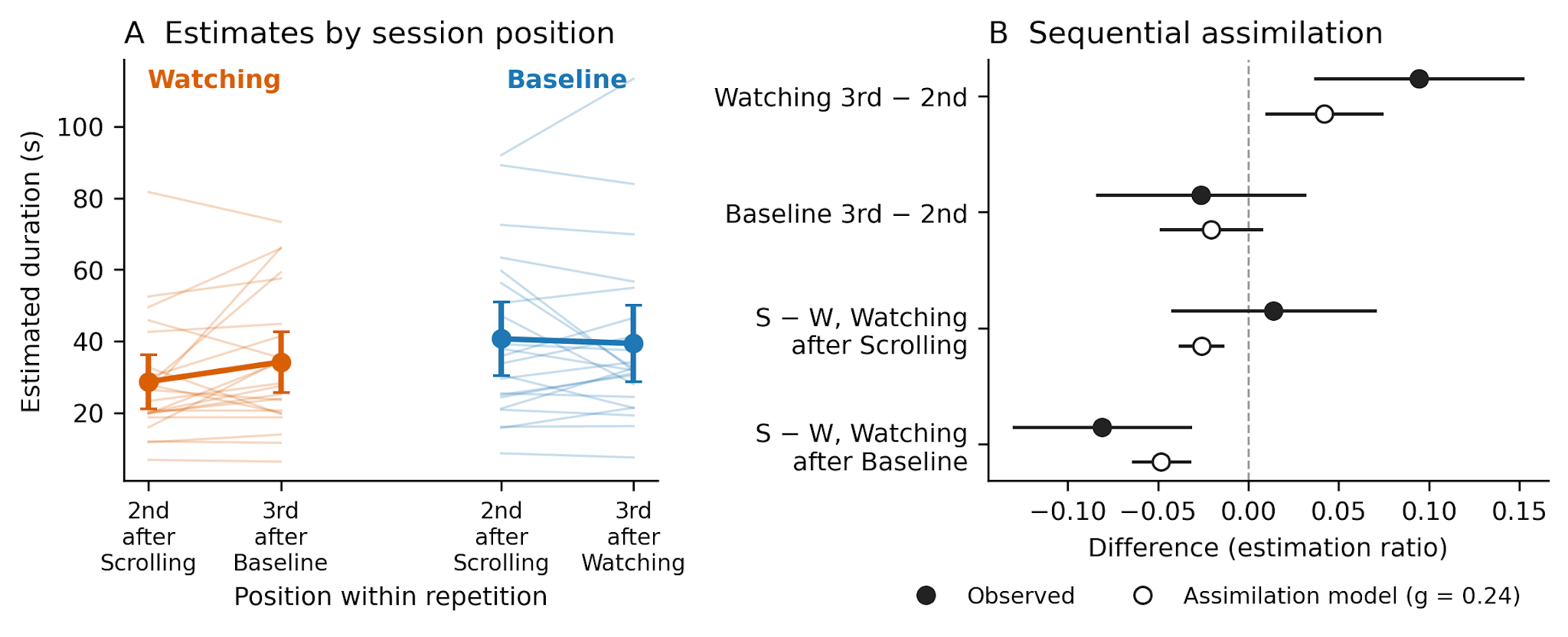

**Figure S4.1.** *Estimates by session position, and the position and context effects predicted by sequential assimilation.* (A) Watching and Baseline estimates when the condition was second (immediately after Scrolling) or third; participant means, lines, and 95% CIs (*n* = 22). (B) Position and context effects in the estimation ratio, observed (filled) and predicted by the sequential-assimilation model (open; each block's ratio drawn toward the preceding block's ratio with weight *g* = 0.24), participant means with 95% CIs; blocks of each session's first sequence, which have no preceding sequence, are excluded (*n* = 22). S − W: Scrolling − Watching, matched on repetition and block length.

Table S4.1. Condition (Watching, Baseline) × Position (second, third) repeated-measures ANOVA on participant-level cell means, *N* = 22. The contrast column gives the paired contrast equivalent to each effect (condition: Baseline − Watching; position: third − second; interaction: Watching shift − Baseline shift).

| **DV** | **Effect** | **F(1, 21)** | **p** | **η_p_²** | **Contrast** | **95% CI** |
| --- | --- | --- | --- | --- | --- | --- |
| Estimate (s) | Condition | 8.92 | .007 | .298 | 8.59 | [2.61, 14.57] |
| Estimate (s) | Position | 5.37 | .031 | .204 | 2.10 | [0.22, 3.99] |
| Estimate (s) | Condition × Position | 1.92 | .180 | .084 | 6.80 | [−3.40, 17.00] |
| Ratio | Condition | 15.42 | < .001 | .423 | 0.138 | [0.065, 0.211] |
| Ratio | Position | 3.80 | .065 | .153 | 0.034 | [−0.002, 0.071] |
| Ratio | Condition × Position | 8.04 | .010 | .277 | 0.121 | [0.032, 0.210] |

The order of conditions explains why Watching shifted and Baseline did not. Watching was preceded either by Scrolling, whose estimates were short, or by Baseline, whose estimates were long; Baseline was preceded by Scrolling or Watching, whose estimates were similar. If each estimate is drawn toward the preceding one, only Watching should move with position. We tested this with a block-level model, in which each block's estimation ratio is drawn toward the preceding block's ratio with weight *g* (participant random intercept; condition, block length, and repetition as factors; 732 blocks with a preceding sequence). Assimilation was reliable, *g* = 0.24 [0.17, 0.31], and the model fitted better than one with position terms instead (AIC −318 vs −282). It reproduces the asymmetry in direction: a Watching shift, almost none for Baseline, and a larger Scrolling − Watching difference after Baseline than after Scrolling (Figure S4.1B). It predicts about half of the observed size of the Watching shift (0.042 vs 0.095 in ratio units), and with assimilation in the model the Condition × Position interaction shrinks from 0.092 to 0.064 (*p* = .052). Carry-over was weakest exactly at condition switches (*g* = 0.07–0.12 for the first block of a sequence), so sequential assimilation explains the direction of the context effect and part of its size, not all of it.

Scrolling's own position can only be examined across repetitions (Figure S4.2A). Scrolling came first in the first repetition too, yet there it did not differ from Watching; the difference appeared from the second repetition on (Condition × Repetition χ²(3) = 2.01, *p* = .570), after overall estimates had risen by 9.8 s [2.4, 17.2], and within a sequence it was largest in the first block (−3.4, −2.8, and −0.8 s). Block-level models over sequences 2–11 recover the difference only with repetition as a factor, because the rise in estimates was a single step rather than a trend. The lag-1 coefficient (−1.49 s) is a lower bound on the increment, because successive blocks share a participant and a condition sequence. From the second repetition on, every Scrolling sequence followed the break and eye-tracker recalibration that closed the preceding repetition, a confound these analyses cannot remove. Baseline − Scrolling was present in every repetition and at both of Baseline's positions (10.7 and 11.3 s); Baseline − Watching was smaller, with an interval including zero, in repetition 2 and when Watching came third, after Baseline (5.2 s [−3.1, 13.5]; Figure S4.2B).

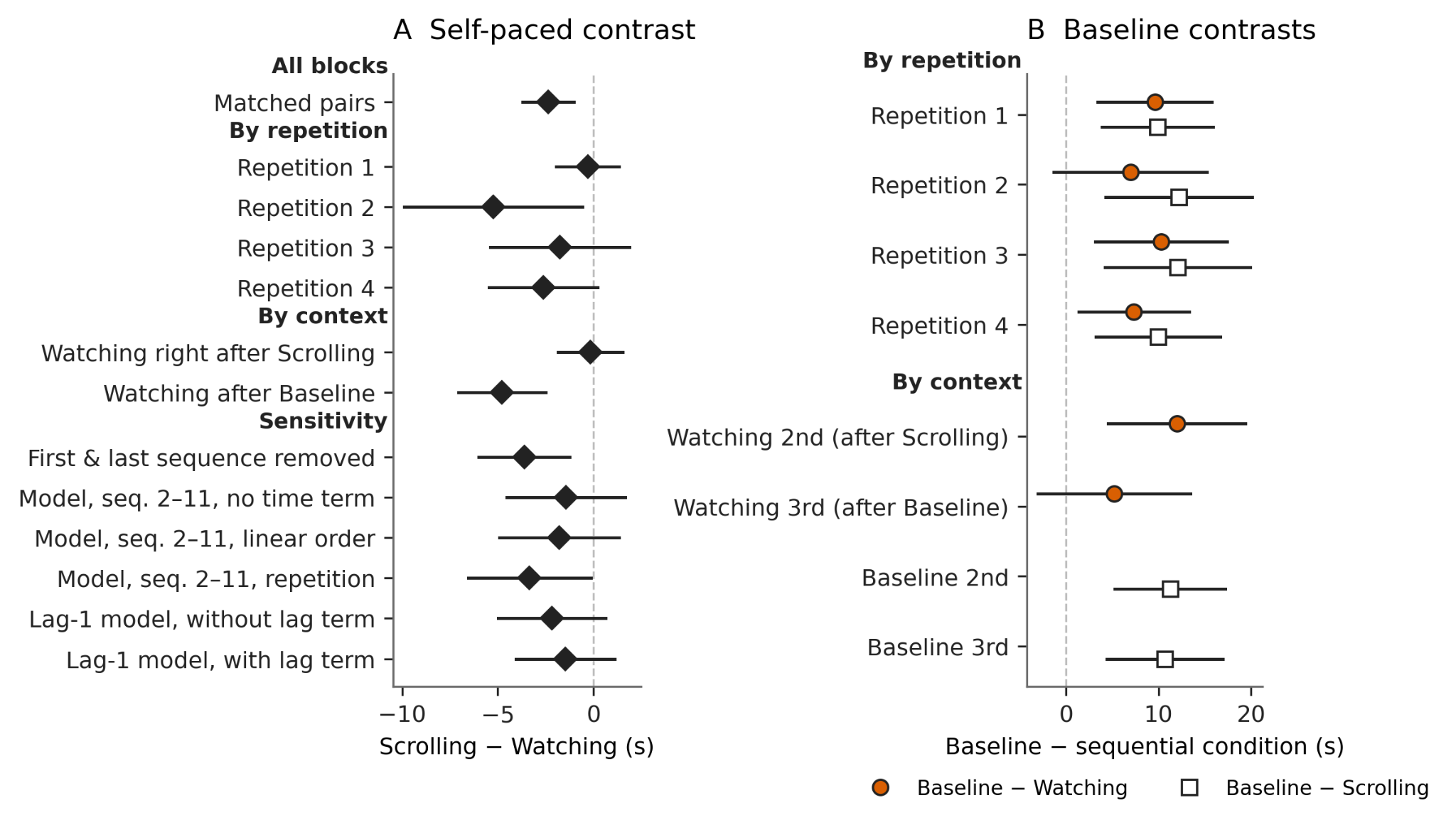

**Figure S4.2.** *Scrolling − Watching and Baseline contrasts across repetitions, contexts, and models (raw estimate).* (A) Scrolling − Watching difference in all matched pairs, within each repetition, by the context in which Watching occurred, after removing the first and last condition sequence, in block-level mixed models over sequences 2–11 (participant random intercept and random slopes for block length) with either no time term, a linear sequence-order term, or repetition as a factor, and in lag-1 models without and with the preceding block's estimate. (B) Baseline − Watching (filled) and Baseline − Scrolling (open) within each repetition and by the position of the block named in each row. Ratio models follow the estimate models (two did not converge). All outputs are in the project repository.

Participants differed widely in their estimation scale (between-participant *SD*s of the condition means 16–23 s) but kept that scale across conditions (within-participant *r* = .98 for Scrolling and Watching and .81–.82 for the Baseline contrasts), so the paired contrasts are precise (*SD* of the paired differences 3.1 s for Scrolling − Watching and 13.2–13.7 s for the Baseline contrasts). Retaining the participant excluded for misunderstanding the task (*N* = 24), or all three excluded participants with complete data (*N* = 26), leaves every contrast in the same direction and of similar size (Figure S4.3). Both Baseline contrasts keep their Holm-corrected significance in every sample (all *p*_Holm_ ≤ .004). The Scrolling − Watching estimate contrast remains significant (*p*_Holm_ = .008 and .044), but its ratio contrast falls short (*p*_Holm_ = .064 and .080).

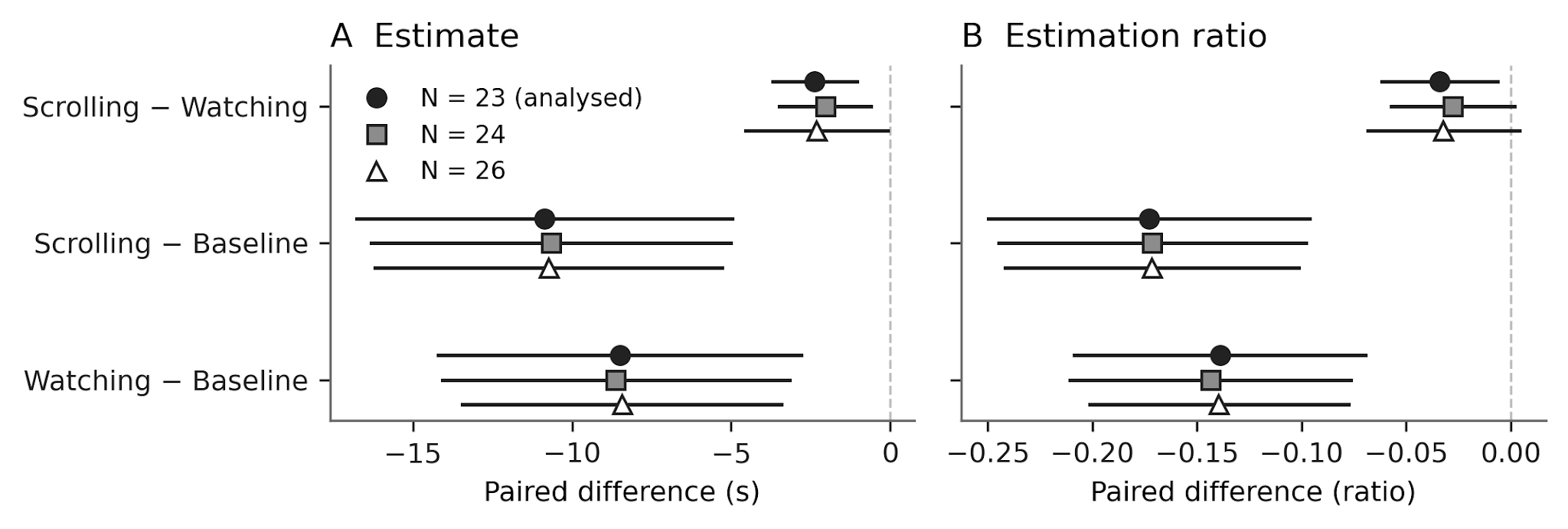

**Figure S4.3.** *Paired contrasts in the analyzed sample and two exclusion-sensitivity samples.* (A) Raw estimate. (B) Estimation ratio. *N* = 23 is the analyzed sample; *N* = 24 adds the participant excluded for misunderstanding the task; *N* = 26 adds all three excluded participants with complete data.

### S5. Picture-locked ERP checks

Baseline-corrected amplitudes at the posterior-occipital cluster were larger under Scrolling than under Watching in all three windows, but none of the differences remained without baseline correction (Figure S5.1), so the effect tracks a pre-onset posterior offset between self-triggered and externally triggered picture streams rather than stronger processing of each picture. Separately, the P2-like positivity reached half its peak earlier under Scrolling (135.3 vs 141.1 ms; Δ = −5.8 ms, *t*(22) = −3.02, *p* = .006, *d*_z_ = −0.63). A cluster-based permutation test over the posterior cluster from 0 to 1000 ms (5000 permutations) found no significant cluster, so the difference is confined to the targeted windows and the rise latency.

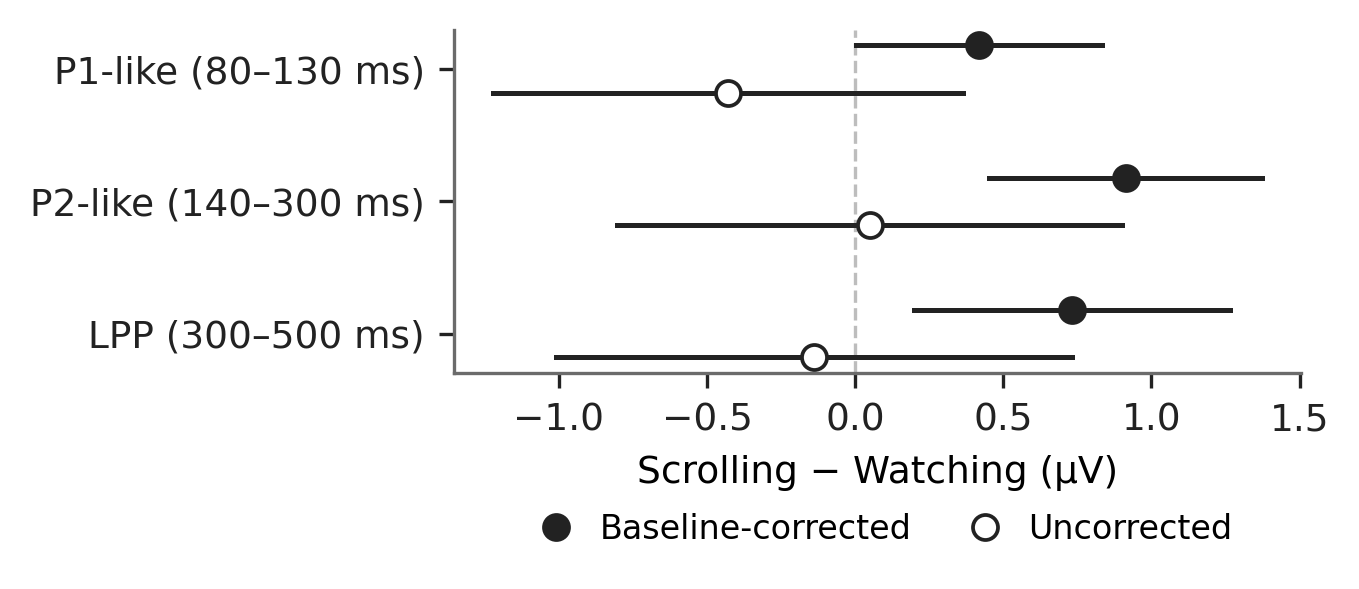

**Figure S5.1.** *Scrolling − Watching picture-locked amplitude by window, with and without pre-stimulus baseline correction.* Posterior-occipital cluster (PO3, PO4, POz, PO7, PO8, O1, Oz, O2). Baseline-corrected: P1-like *p* = .048, P2-like *p* < .001, LPP *p* = .010; uncorrected: *p* = .274, .906, and .743.

### S6. Fixation-locked FRP checks

The planned fixation-locked measure is the occipital lambda response (60–120 ms). An exploratory grid of three clusters and four windows (Figure S6.1; uncorrected) shows why the main text restricts its claim to lambda: at the occipital cluster, the later windows separated both sequential-picture conditions from Baseline (all *p* < .001) but showed no Scrolling-over-Watching enhancement, with a small difference in the opposite direction for P2 (*p* = .070). The occipital contrasts held in single-fixation mixed models with crossed random intercepts for participant and image (69,133 fixation epochs, 374 images; Figure S6.2A). Adding incoming saccade amplitude (0.23 µV per degree, *t* = 36.2) reduced the Scrolling − Watching lambda coefficient by 72%, and condition no longer improved the model (χ²(1) = 0.70, *p* = .401; Figure S6.2B). The per-fixation lambda difference therefore reflects the larger saccades of self-paced viewing.

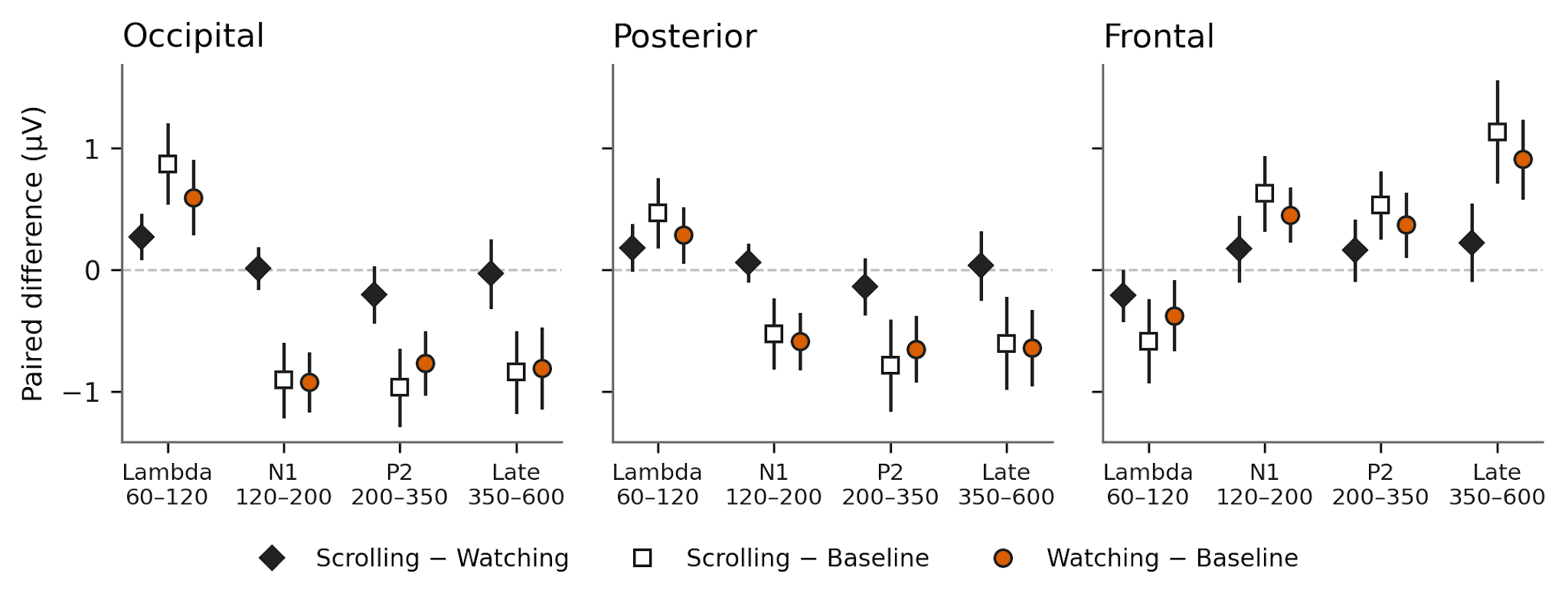

**Figure S6.1.** *Exploratory fixation-locked contrasts by cluster and window (ms).* Participant-level paired differences with 95% CIs; *p* values are uncorrected and the grid is exploratory except for occipital lambda.

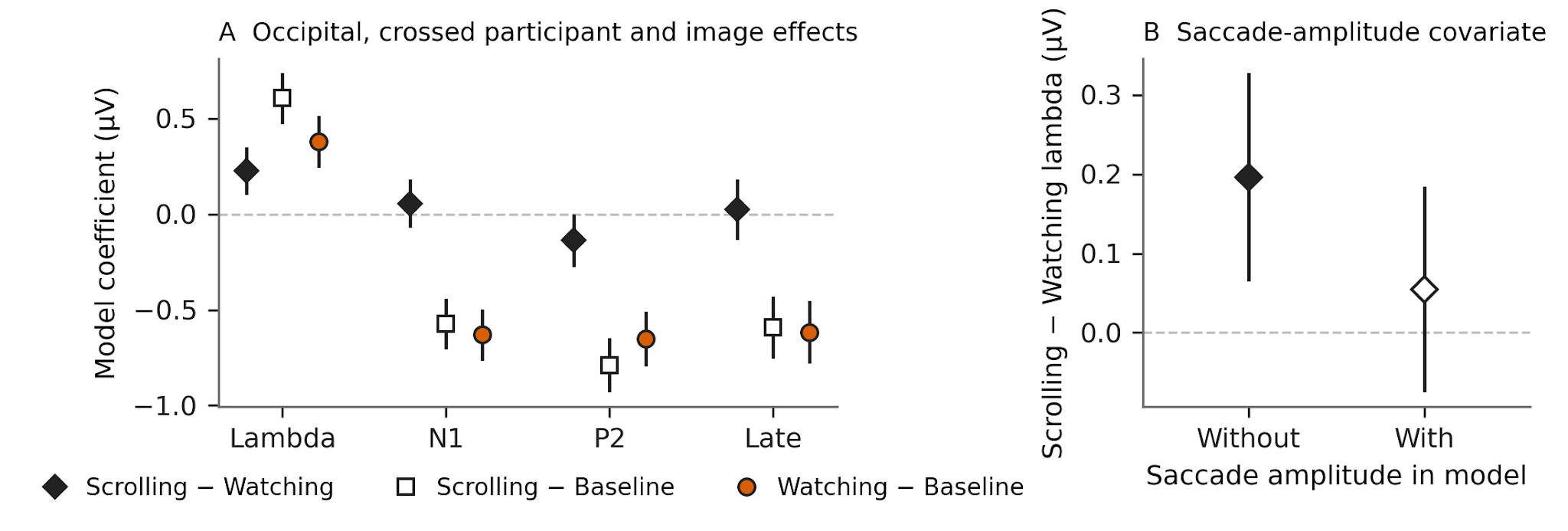

**Figure S6.2.** *Single-fixation mixed models of occipital fixation-locked amplitude.* (A) Condition coefficients by window with participant and image random intercepts. (B) Scrolling − Watching lambda coefficient without and with incoming saccade amplitude as a covariate, in the subset of fixations with a reconstructable preceding saccade (35,481 Scrolling and Watching epochs).
